# Identification of Transient Intermediates During Spliceosome Activation by Single Molecule Fluorescence Microscopy

**DOI:** 10.1101/2022.04.23.488636

**Authors:** Xingyang Fu, Harpreet Kaur, Margaret L. Rodgers, Eric J. Montemayor, Samuel E. Butcher, Aaron A. Hoskins

**Author notes:** Margaret L Rodgers, T.C. Jenkins Department of Biophysics, Johns Hopkins University, Baltimore, MD, 21218, USA.

## Abstract

Spliceosome activation is the process of creating the catalytic site for RNA splicing and occurs *de novo* on each intron following spliceosome assembly. Dozens of factors bind to or are released from the activating spliceosome including the Lsm2-8 heteroheptameric ring that binds the U6 small nuclear RNA (snRNA) 3’-end. Lsm2-8 must be released to permit active site stabilization by the Prp19-containing complex (NineTeen Complex, NTC); however, little is known about the temporal order of events and dynamic interactions that lead up to and follow Lsm2-8 release. We have used colocalization single molecule spectroscopy (CoSMoS) to visualize Lsm2-8 dynamics during activation of yeast spliceosomes. Lsm2-8 is recruited as a component of the tri-snRNP and is released after integration of the Prp19-containing complex (NineTeen Complex, NTC). Despite Lsm2-8 and the NTC being mutually exclusive in existing cryo-EM structures of yeast B complex spliceosomes, we identify a transient intermediate containing both 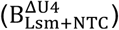 and provide a kinetic framework for its formation and transformation during activation. Prior to 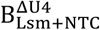 assembly, the NTC rapidly and reversibly samples the spliceosome suggesting a mechanism for preventing NTC sequestration by defective spliceosomes that fail to properly activate. In complementary ensemble assays, we show that a base-pairing dependent ternary complex can form between Lsm2-8 and U2 and U6 helix II RNAs. Together our data suggest a Hfq-like function for Lsm2-8 in maintaining U2/U6 helix II integrity before it can be transferred to the NTC by transient formation of the 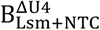 spliceosome.

**Significance Statement:** The spliceosome active site is created *de novo* during activation and involves numerous conformational and compositional changes. Here, we define a kinetic pathway for yeast spliceosome activation using single molecule fluorescence that includes transient intermediates not previously identified. Real-time measurements allow us to uncover rapid, reversible sampling interactions of the NineTeen Complex (NTC) that may prevent its accumulation on defective spliceosomes. By analogy with bacterial Hfq, we propose that the homologous Lsm2-8 proteins stabilize U2/U6 helix II during activation before the helix is transferred to the NTC in a short-lived spliceosome containing both Lsm2-8 and the NTC. Our data demonstrate how single molecule studies of activation can reveal kinetically-competent intermediates and complement cryo-EM studies of stalled or inhibited complexes.

## INTRODUCTION

Pre-mRNA splicing is an essential step in eukaryotic gene expression. Disruption of the splicing process on a molecular level can lead to human diseases including retinitis pigmentosa, cancers, and amyotrophic lateral sclerosis (1, 2). Molecular mechanisms of splicing have been studied for decades *in vitro* and *in vivo* by a combination of techniques. The overall splicing process is evolutionarily conserved and involves formation of numerous intermediate spliceosome complexes (**Figs. 1A, S1**) (3-6). The building blocks of spliceosomes include five small nuclear ribonucleoproteins (U1, U2, U4, U5, and U6 snRNPs), each made of one snRNA and several protein factors. These snRNPs undergo assembly on the substrate, activation to form the catalytic site, catalysis of two sequential transesterification reactions, and disassembly/recycling. The NTC and NTC-related factors (NTR) are protein-only subunits that are necessary for catalysis and join the spliceosome during activation (7, 8). A successful splicing process is achieved by assembly and disassembly of intricate interaction networks involving 5 snRNAs and over 170 protein factors in humans and ∼90 in *S. cerevisiae* (yeast) (5, 9).

**Figure 1.**
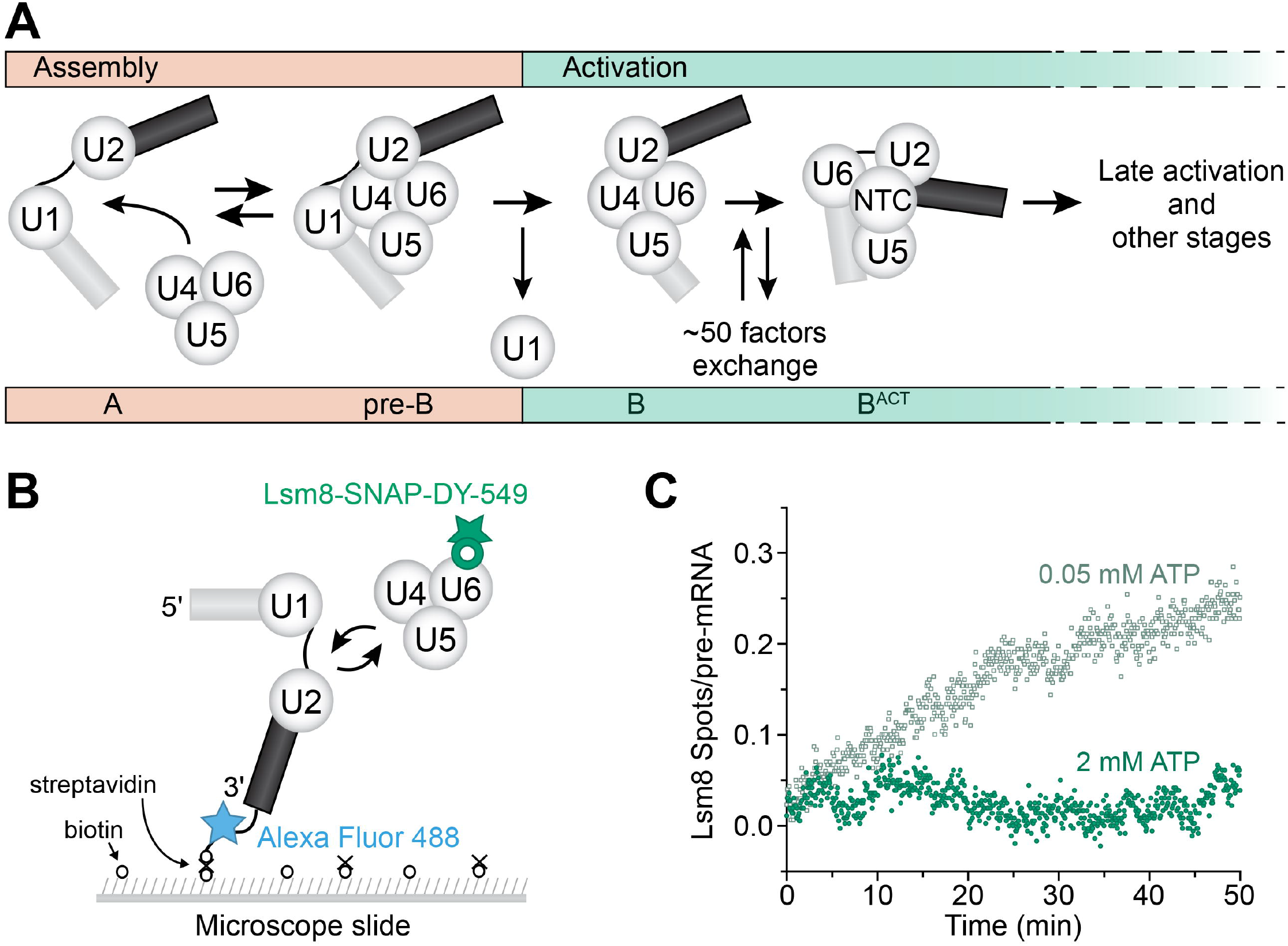
2-Color CoSMoS Assay to Study Lsm8 Dynamics During Spliceosome Activation. (**A**) The U4/U6.U5 tri-snRNP reversibly interacts with the spliceosome A complex consisting of U1 and U2 snRNPs associated with pre-mRNA. Once the pre-B complex is formed, ATP-dependent activation can proceed by first ejecting the U1 snRNP to form the B complex. This is then followed by exchange of ∼50 protein and snRNA factors to form the activated B complex (B^ACT^ spliceosome). During these steps, the U4 snRNP and Lsm2-8 proteins are released, and the NTC joins the spliceosome. Subsequent steps then lead to formation of the spliceosome active site and splicing. (**B**) Schematic of a 2-color CoSMoS experiment for observing Lsm ring binding dynamics. The U4/U6.U5 tri-snRNP contains a DY-549 fluorophore attached to the U6 Lsm ring component Lsm8 with a SNAP tag. The pre-mRNA was immobilized to the slide surface and contained an Alexa 488 fluorophore. (**C**) Time record of the number of Lsm8 fluorescent spots that appeared on the surface relative to the number of surface-tethered pre-mRNA molecules at 0.05 mM (grey) and 2 mM ATP (green). At 0.05 mM ATP, spliceosome pre-B complexes can form but activation cannot proceed. At 2 mM ATP, spliceosome activation and splicing can proceed.

The Sm and Lsm2-8 protein complexes are two important factors that contribute to pre-mRNA splicing by chaperoning snRNAs during snRNP biogenesis and forming interactions with other splicing factors (10-13). Sm and Lsm2-8 proteins in the spliceosome form heteroheptameric rings with small central channels for RNA binding (14). Individual Sm or Lsm proteins share a common fold with the Hfq protein, which forms homohexamers and regulates sRNA/mRNA annealing in bacteria (15). The Lsm2-8 complex preferentially binds to the post-transcriptionally processed 3’ end of U6 snRNA (16) and is present in the U6 snRNP and U4/U6.U5 tri-snRNP. Lsm2-8 remains bound to U6 during spliceosome assembly but is released from yeast spliceosomes during activation in a process that requires the NTC (17). For the yeast splicing machinery, it is not known if Lsm2-8 release occurs before, after, or concertedly with NTC association; however, recent cryo-EM structures of human spliceosomes show that spliceosome complexes containing both Lsm2-8 and a subset of NTC proteins can form (18). Formation of the spliceosome active site has been proposed to be directed by a series of mutually exclusive interactions in which Lsm2-8 and the NTC play critical roles (18). Therefore, knowledge of the kinetic pathways for Lsm2-8 release and NTC recruitment is essential for understanding activation of spliceosomes.

Spliceosome activation is complex and involves dozens of coordinated compositional and conformational changes. For example, the transition from the yeast B to B^ACT^ spliceosome involves the exchange of around 50 different factors (5) (**Figs. 1A, S1**), making it intrinsically difficult to study. While cryo-EM has captured snapshots of human spliceosome activation intermediates (18) and has provided key insights on the mechanism of splicing (13), it is difficult to derive kinetic pathways from these data. Previously, our lab has used Colocalization Single Molecule Spectroscopy (CoSMoS) to study the activation pathways of single spliceosomes by monitoring tri-snRNP binding, U4 snRNP protein release, and NTC recruitment (19). Single molecule techniques like CoSMoS can resolve heterogenous subpopulations, reveal new intermediates, and identify the temporal order of events occurring in complex, unsynchronized assembly processes (19-21). Prior CoSMoS studies of activation showed that while tri-snRNP binding was reversible, U4 snRNP release during activation is an irreversible step and that NTC association occurs predominantly after loss of U4 proteins (19). Whether or not Lsm2-8 release represents another irreversible step and how this is kinetically coupled with NTC recruitment are not known.

Here, we used CoSMoS to study Lsm2-8 dynamics during activation. Our results show that Lsm2-8, as expected, associates with the spliceosome as part of the tri-snRNP. Unexpectedly, Lsm2-8 release occurs after association of the NTC suggesting formation of B complex spliceosome lacking the U4 snRNP but containing both the NTC and Lsm2-8 ring 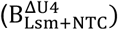. Formation of this complex is often preceded by the NTC reversibly sampling the spliceosome, and stable NTC binding is followed by essentially irreversible release of Lsm2-8. Combined with *in vitro* biochemical data for ternary complex formation between Lsm2-8, U2, and U6 RNAs, we propose a kinetic model for spliceosome activation involving formation of at least two transient intermediates of unknown structure including the 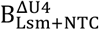 complex. We propose that release of Lsm2-8 from the 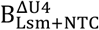 complex may help to maintain U2/U6 helix II integrity during activation until the duplex can be bound by NTC proteins.

## RESULTS

### ATP-Dependent Accumulation of Lsm8 Complexes on Single Pre-mRNAs

To watch single molecules of Lsm8 assemble with and release from spliceosomes, we first genetically encoded a c-terminal fast SNAP tag on the Lsm8 protein. Lsm8 is essential for yeast growth in the absence of U6 snRNA overexpression (22), and it is the only member of the Lsm2-8 complex not found in Lsm1-7 complexes involved in mRNA decay. Lsm8-SNAP strains are viable, indicating that the tagged protein is functional for splicing *in vivo*. Lsm8-SNAP can be readily labeled with benzylguanine-fluorophores in yeast whole cell extracts (WCEs; **Fig. S2**) and these WCEs possessed high *in vitro* splicing activities (**Fig. S3**).

To observe Lsm8 dynamics, we labeled Lsm8 with SNAP-DY-549 [a 532nm (green) laser-excitable fluorophore] in WCE and added the WCE to passivated glass slides containing RP51A pre-mRNAs labeled with Alexa Fluor 488 [a 488nm (blue) laser-excitable fluorophore] and immobilized with biotin-streptavidin (**Fig. 1B**). Experiments were carried out at two concentrations of ATP: 0.05 mM ATP, which permits spliceosome assembly but not activation or Lsm release, and 2 mM ATP, which permits activation, Lsm release, and splicing (7). Using a custom-built CoSMoS microscope (23), we observed fixed fluorescent RNA spots upon 488 nm laser excitation and dynamic spots originating from Lsm8-SNAP upon 532 nm laser excitation.

The Lsm8-SNAP binding and unbinding events were dependent on ATP concentration. At 0.05 mM ATP, Lsm8-SNAP fluorescent spots persisted, and this resulted in their surface accumulation over time (**Fig. 1C**). In contrast, at 2 mM ATP, Lsm8-SNAP fluorescent spots appeared more briefly, and this resulted in little surface accumulation (**Fig. 1C**). The surface accumulation data strongly resemble those previously obtained for the U4 snRNP under these ATP conditions (19) and are consistent with assembly of Lsm8-SNAP-containing spliceosomes that cannot undergo activation and retain Lsm8-SNAP at 0.05 mM ATP. At 2 mM ATP, Lsm8-SNAP-containing spliceosomes can both assemble and activate, resulting in Lsm8-SNAP release and transient association with the surface-tethered RNAs.

### The Lsm Ring is Released after U4 snRNP Dissociation

To identify Lsm8-SNAP proteins associated with tri-snRNPs and correlate Lsm8 release with spliceosome activation, we carried out 3-color CoSMoS assays using extracts containing Lsm8-SNAP labeled with DY-549 fluorophores, the U4 snRNP proteins Prp3 and Prp4 both labeled with the DHFR tag and Cy5-TMP [632nm (red) laser-excitable] fluorophores, and Alexa Fluor 488 labeled pre-mRNAs (**Fig. 2A**). We have previously used CoSMoS to show that Prp3 and Prp4 are both lost during spliceosome activation and that their release is irreversible (19). In agreement with these previous experiments, Prp3/Prp4 fluorescence spots accumulated to a greater extent on a surface containing immobilized pre-mRNAs at 0.05 mM ATP than at 2 mM ATP (**Fig. S4**). In these 3-color experiments, the apparent trends of surface accumulation of Lsm8-SNAP and Prp3/Prp4-DHFR were quite similar at both ATP concentrations (**Figs. S4C, D**). Since Lsm8, Prp3, and Prp4 are all components of the tri-snRNP, the similarity likely originates from tri-snRNP binding events.

**Figure 2.**
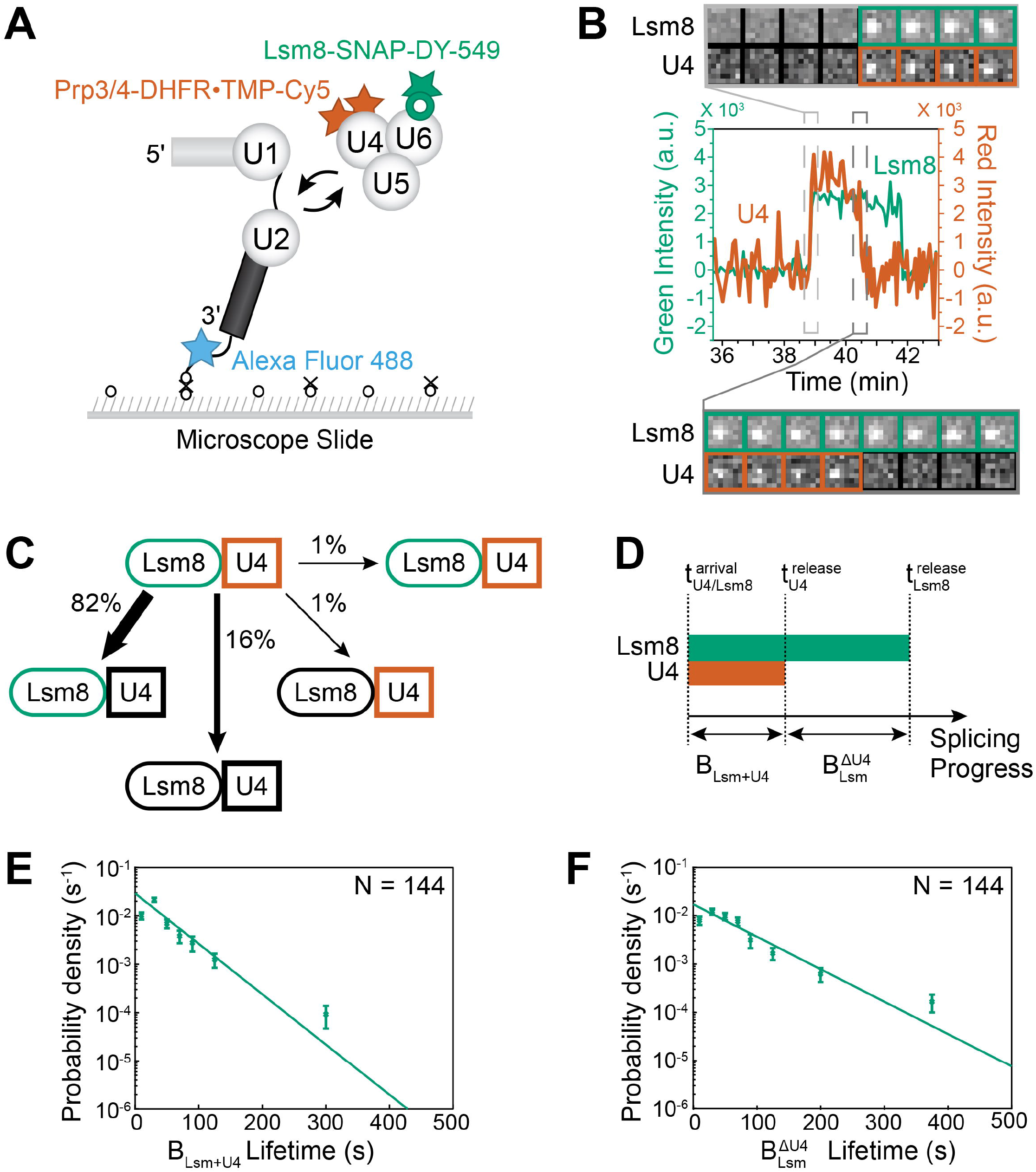
3-color CoSMoS observation of U4 snRNP and Lsm ring binding dynamics during activation. (**A**) Schematic of a 3-color experiment in which U4 was labeled with Cy5-TMP fluorophores, the Lsm ring was labeled with a DY-549 fluorophore, and the surface-tethered pre-mRNA was labeled with an Alex Fluor 488 fluorophore. (**B**) Segment of a representative time record showing peaks in fluorescence intensity corresponding to colocalization of U4 (red, thick line) and Lsm8 proteins (green, thin line) with the same individual pre-mRNA molecule. The light grey dashed rectangle marks an example of the simultaneous appearance of U4 and Lsm8 spots; galleries show consecutive images taken from the indicated part of the recording showing that spot appearance is simultaneous. The darker grey dashed rectangle marks an example of the ordered disappearance of U4 then Lsm spots; galleries show consecutive images taken from the indicated part of the recording showing that loss of the U4 signal precedes loss of the Lsm8 signal. (**C**) Routes for loss of either the U4 or Lsm fluorescent spots at 2 mM ATP for *N*=176 pairs of overlapping events. Green and red shapes represent observation of fluorescence from the corresponding DY-549 or Cy5 fluorophores on Lsm or U4, respectively; grey shapes represent the absence of fluorescence. Percentages represent the fraction of U4/Lsm complexes in which fluorescence disappeared by the indicated pathway; more prevalent pathways are emphasized with thicker arrows. (**D**) Schematic showing the relationship between the lifetimes of the B_Lsm+U4_ and 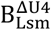 complexes to the measured arrival and release times for U4 snRNP and Lsm8 proteins. (**E, F**) Probability density histograms of B_Lsm+U4_ (panel E, 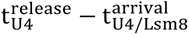) and 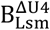 (panel F, 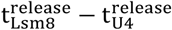) complex lifetimes obtained from the subset of events (*N*) showing simultaneous arrival of U4 and Lsm spots followed by ordered loss of the U4 and then Lsm signals. Lines represent fits of the lifetime distributions with equations containing single exponential terms that yielded the parameters reported in **Table S1**. Error bars (SD) were calculated for each point as described in the Methods.

We analyzed individual fluorescence time trajectories to identify events at pre-mRNA locations in which Lsm8-SNAP and Prp3/Prp4-DHFR fluorescence colocalized (**Fig. 2B, S5**). Approximately 28% of Lsm8-SNAP or Prp3/Prp4-DHFR events at 2 mM ATP colocalized with one another at some point during the event lifetime (201 out of 723 Lsm8-SNAP events or 685 Prp3/Prp4-DHFR events). Since these proteins are expected to be stoichiometric with one another in the tri-snRNP, this suggests either incomplete fluorophore labeling or potential tri-snRNP heterogeneity. We restricted our analysis just to the subset of colocalized events. Of these, the majority (88%) showed simultaneous arrival of Lsm8-SNAP and Prp3/Prp4-DHFR fluorescent spots. This indicates that the predominant recruitment pathway for Lsm2-8 is concurrent with the U4 snRNP for the colocalized signals. This agrees with biochemical and structural data for the tri-snRNP indicating simultaneous presence of these factors (24-26).

For this subset of Lsm8-SNAP and Prp3/Prp4-DHFR events, we then analyzed the order in which their signals disappeared from the immobilized RNAs. At 2 mM ATP, the predominant release pathway (82%, 144/176 events) showed loss of the Lsm8-SNAP signal after loss of the Prp3/Prp4-DHFR signals (**Fig. 2B, C**). The losses of the Lsm8-SNAP signals at 2 mM ATP were unlikely to be due entirely to photobleaching since we readily observed much longer-lived signals at 0.05 mM ATP (∼8-fold longer fitted longest-lived lifetime, **Table S1**). Thus, we interpret these observations as U4 snRNP dissociation occurring prior to Lsm2-8 release during yeast spliceosome activation. This is in agreement with a proposed activation pathway for the human spliceosome deduced from cryo-EM structures of pre-B^ACT^ complexes (18).

The single molecule data support the existence of at least two spliceosomes with distinct compositions: a B_Lsm+U4_ complex that contains Lsm8, Prp3, and Prp4 and a 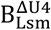 complex that contains Lsm8 but neither Prp3 nor Prp4. The B_Lsm+U4_ complex likely represents the yeast pre-B or B complex spliceosome that has been previously biochemically and structurally characterized (25, 27). In these complexes, the tri-snRNP has associated with the U1- and U2-containing A complex but activation has not yet proceeded to the point of U4 snRNP release. The 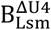 intermediate is characterized by its formation from the B_LSM+U4_ complex, the absence of U4 snRNP proteins, and the presence of Lsm8. The 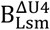 complex, therefore, represents an activation intermediate formed after release of the U4 snRNP but from which the Lsm2-8 ring has not yet been released. It should be noted that in these experiments we cannot discriminate between 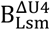 complexes that contain the NTC and those that do not (see the following section). Though 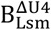 has not been previously identified in yeast, a cryo-EM structure of a human spliceosome containing a similar composition (pre-B^ACT1^) has been obtained in the presence of an activation inhibitor (18).

To provide further insight into the B_Lsm+U4_ and 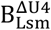 complexes, we analyzed the arrival and departure times for each colocalized Lsm8-SNAP and Prp3/Prp4-DHFR event. The lifetimes of the B_Lsm+U4_ complexes were determined by subtracting the arrival times of the colocalized signals from the U4 release times 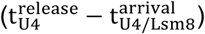, and the lifetimes of the 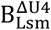 complexes were found by subtracting the Prp3/Prp4-DHFR release times from the release times of the Lsm8-SNAP signals 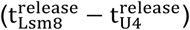 (**Fig. 2D**). The resulting unbinned distributions of bound dwell times were then fit using maximum likelihood methods to exponential-based functions (28).

In both cases, the distributions could be best fit to functions containing single exponential terms with characteristic lifetimes of 41.7±5.3 and 64.7±6.9 s for B_Lsm+U4_ and 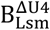 complexes, respectively (**Fig. 2E, F; Table S1**). The lifetime of the B_Lsm+U4_ complex is very similar to that previously measured for U4 snRNP proteins (∼ 34 s) under activation conditions (19) and likely represents the average residence time for the U4 snRNP in pre-B/B complex spliceosomes. Lsm8 release does not happen immediately after loss of U4 proteins. Instead, Lsm8 remains bound to the spliceosomes for ∼ 1 min as represented by the lifetime of the 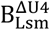 complex. The lifetimes for both B_Lsm+U4_ and 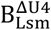 complexes are smaller relative to the *in vitro* timescale of RP51A intron loss, which occurs over tens of minutes (29). This indicates that both complexes are kinetically competent intermediates for activation. Finally, since we could measure the lifetimes of complexes formed in succession from the same molecules, we analyzed whether the lifetimes of the two complexes correlated with one another. In other words, we asked if rapid release of U4 was predictive of subsequent slow or rapid release of Lsm8 from the same molecular complex. Analysis of the paired B_Lsm+U4_ and 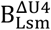 lifetimes did not reveal a strong correlation between the two (**Fig. S6A**). This suggests that U4 snRNP release kinetics are to some degree decoupled from the release of the Lsm ring with the rate of the former not predictive of the latter.

### The NTC Joins the Spliceosome Before Lsm2-8 Release

We next investigated the timing of Lsm8-SNAP release relative to NTC recruitment during activation. Previous work from our lab showed that the NTC associates after U4 snRNP release (19). Since Lsm2-8 release also occurs after U4 snRNP release, it is possible that the NTC associates while the Lsm2-8 ring is still present, concertedly with Lsm2-8 departure, or afterwards. To discriminate between these possibilities, we carried out 3-color CoSMoS assays using extracts containing Lsm8-SNAP labeled with DY-549, NTC proteins Cef1 and Syf1 both labeled with the DHFR tag and Cy5-TMP, and Alexa Fluor 488-labeled pre-mRNAs (**Figs. 3A, S3**).

**Figure 3.**
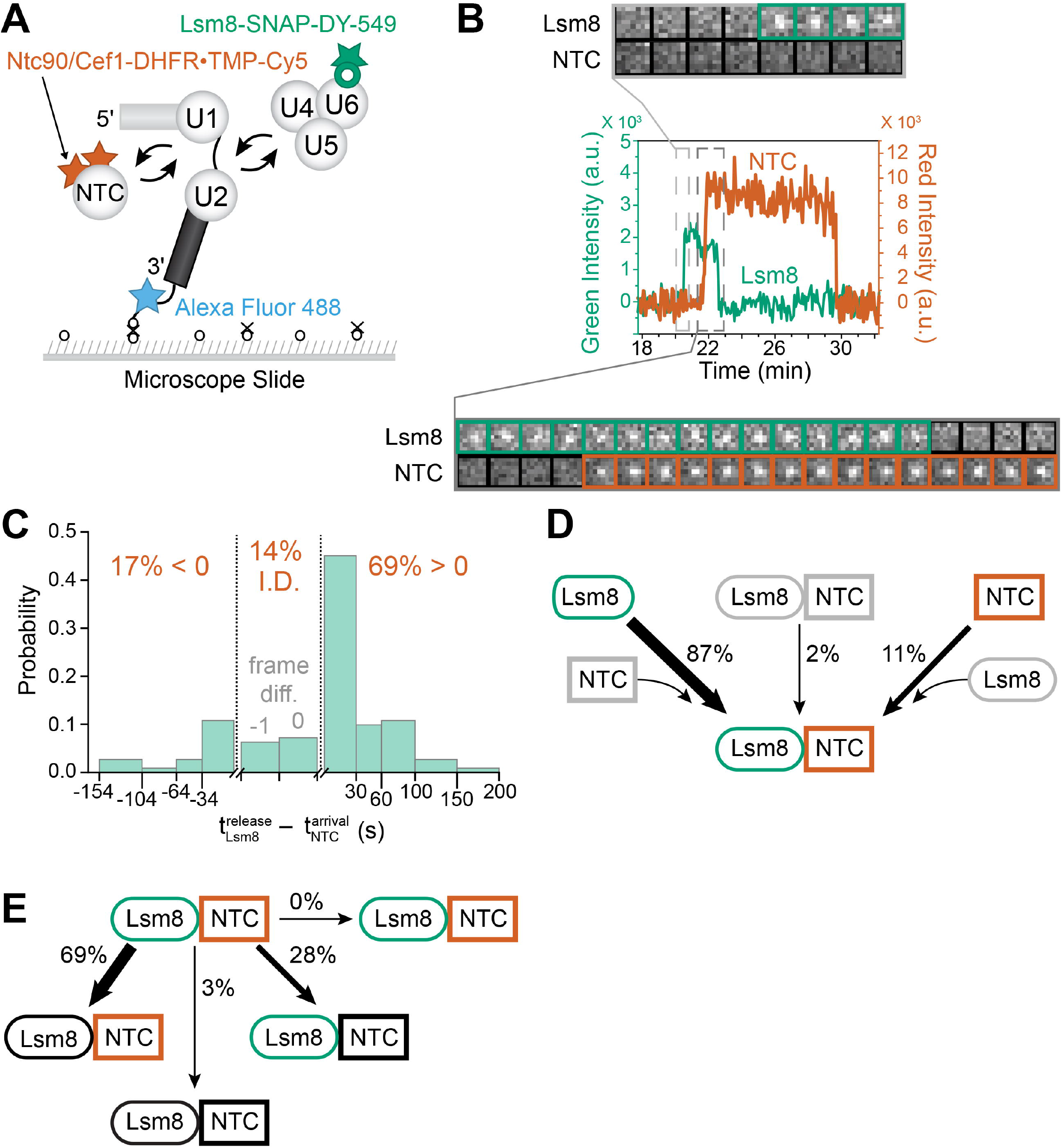
Three-color CoSMoS observation of NTC and Lsm ring binding dynamics during activation. (**A**) Schematic of a 3-color experiment in which the NTC was labeled with Cy5, the Lsm ring was labeled with a DY-549, and the surface-tethered pre-mRNA was labeled with Alexa Fluor 488. (**B**) Segment of a representative time record showing peaks in fluorescence intensity corresponding to colocalization of NTC (red, thick line) and Lsm proteins (green, thin line) with the same individual pre-mRNA molecule. The light grey dashed rectangle marks an example of the ordered appearance of NTC and Lsm8 spots; galleries show consecutive images taken from that part of the recording showing that spot appearance is not simultaneous. The dark grey dashed rectangle marks an example of simultaneous occupancy of the pre-mRNA by the NTC and Lsm ring followed by the ordered disappearance of Lsm8 and then NTC spots; galleries show consecutive images taken from the indicated part of the recording showing overlap of the Lsm8 and NTC signals. (**C**) Probability density histogram showing the delay between NTC arrival and Lsm8 release. Most often (69% of *N*= 293 total events), the NTC arrived before release of the Lsm ring (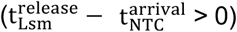> 0). In 14% of cases, the exact order of events was indeterminate (I.D.) as described in the text. (**D**) Routes for the appearance of Lsm8 and NTC fluorescent spots at 2 mM ATP for *N*=157 pairs of overlapping events. (**E**) Routes for loss of either Lsm or NTC fluorescent spots at 2 mM ATP for *N*=136 pairs of overlapping events in which the Lsm8 spot appearance preceded arrival of the NTC.

Analysis of the fluorescence trajectories shows that Lsm8 signals frequently appeared before those from the NTC on single pre-mRNA molecules (**Figs. 3B, S7**). This agrees with the ordered addition of the tri-snRNP and the NTC to spliceosomes (17, 20). Most often the Lsm8 and Cef1/Syf1 signals were observed simultaneously on the same pre-mRNA molecules. This suggests that the NTC associates with the spliceosome during activation before Lsm2-8 release. To determine the frequency of these type of events, we identified all pairs of Lsm8 and Cef1/Syf1 signals and determined the distribution of binding behaviors by calculating the difference between the Lsm8 release and NTC recruitment times 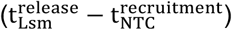 (**Fig. 3C**). Within the distribution, events in which the NTC binds before Lsm8 release result in positive values and events in which the NTC binds after Lsm8 release result in negative values. The order of Lsm8 release and NTC binding cannot be determined (indeterminate, I.D.) for events in which the last frame with a Lsm8 signal is the same as the first frame with a NTC signal 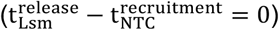 or for cases where the last frame with a Lsm8 signal is immediately followed by the first frame with a NTC. In this latter case, colocalization cannot be judged during the time lapse interval between frames. Of the analyzed event pairs, the majority (∼69%) have 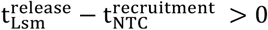, indicating the predominant pathway for NTC protein binding is while Lsm8 is still present. In only ∼17% of event pairs did loss of the Lsm8 signal precede NTC binding. Among the 157 pairs of events showing simultaneous presence of Lsm8 and NTC proteins on the pre-mRNA molecules, 87% showed that Lsm2-8 is recruited prior to NTC arrival (**Fig. 3D**). This is consistent with tri-snRNP recruitment occurring before the integration of the NTC complex (19) and with the NTC arriving before Lsm2-8 release.

It has been proposed that for human spliceosomes undergoing activation, NTC proteins bind sequentially (18). A NTC subcomplex containing CDC5L (the human homolog of Cef1) binds first to form the pre-B^ACT1^ spliceosome. A different subcomplex containing Syf1 (the intron binding complex, IBC) then binds in a second step to form the pre-B^ACT2^ spliceosome. Since in our experiments, Cef1 and Syf1 were both DHFR labeled, we would expect to see a stepwise increase in Cy5-TMP fluorescence intensity during activation if the factors bound sequentially. While we could observe stepwise decreases fluorescence intensity consistent with the presence of both Cef1 and Syf1 followed by loss of the signal from one of the proteins (14 out of 83 NTC events analyzed, **Table S2**), we never observed a stepwise increase in fluorescence consistent with sequential binding. This is unlikely due to sub-stoichiometric occupancy of Cy5-TMP on the labeled proteins, since nearly identical results were previously reported for single molecule studies of Cef1 and Syf1 labeled with the SNAP tag (20). These data suggest that in yeast, Cef1 and Syf1 are recruited to the spliceosome either very rapidly one after another or as part of the same NTC complex.

### NTC Transiently Samples the Spliceosome during Activation

For the subset of molecules with simultaneous occupancy by Lsm8 and Cef1/Syf1, we classified each pair of events according to one of fourteen predicted patterns (**Fig. S8**). While most events showed release of the NTC proteins occurring after Lsm8 release (seq-13 events; **Fig. S8C**), we were surprised to see many events (24% of events), in which the NTC bound and released while the Lsm8 was still present (seq-15 events; **Figs. 4A, S8C**). These NTC events were often short-lived and lasted only a few frames, much shorter than other event types (**Fig. 4B**).

**Figure 4.**
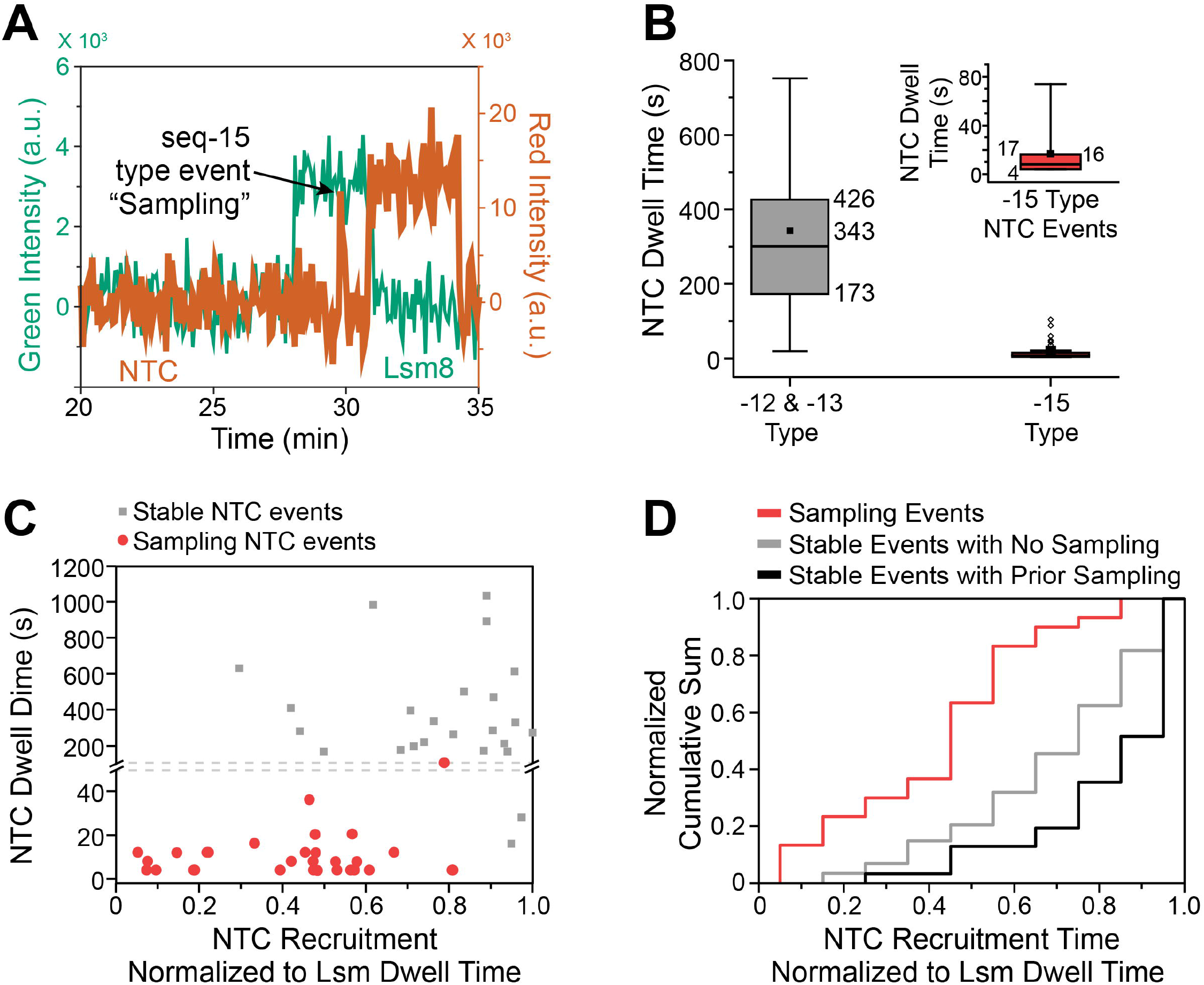
The NTC Frequently Samples the Spliceosome Prior to Stable Binding. **(A**) Example fluorescence trajectory showing a seq-15-type event/NTC sampling (arrow). (**B**) The observed dwell times for sampling events (−15 type) are much shorter than those observed for non-sampling events (−12 and -13 type). The inset shows the -15 type events with an expanded y-axis. Numbers adjacent to the boxes indicate the interquartile range (IQR, 25-75%) and mean. Outliers (hollow circles) are defined as those with dwell times >1.5xIQR. (**C**) Sampling events are more often observed soon after Lsm8 arrival and longer-lived NTC binding events are more often observed closer to the point of Lsm8 release. Fluorescence trajectories were synchronized post-experimentally to the Lsm8 dwell times which were then normalized (time = 0 represents the arrival of the Lsm8 signal and time = 1 represents loss of the Lsm8 signal). The dashed line indicates a break in the y-axis. (**D**) Normalized cumulative sum histogram for NTC binding normalized to Lsm8 dwell times for NTC binding events that do (light and dark green) and do not (blue) include sampling-type events. Long-lived, stable NTC binding events are more likely to occur towards the end of the Lsm8 binding time regardless if a sampling event was observed or not.

It is possible that the short-lived NTC events represent transient interactions with the spliceosome during activation in which stable integration of the NTC is not yet possible (sampling). We predicted that sampling is more likely to occur early in activation since the conformational changes needed for stable NTC binding may not yet be completed. To test this, we synchronized the time of each NTC molecule’s recruitment to the normalized lifetime of the colocalized Lsm8 binding event. A plot of these synchronized NTC recruitment times versus their corresponding NTC dwell times shows that short-lived NTC events are more commonly observed soon after the Lsm8 signal appears (**Fig. 4C**). Longer-lived NTC binding events appear only after ∼50% of the Lsm8 dwell time on the spliceosome has elapsed. These observations are confirmed when we calculate the cumulative sum for NTC binding as a function of the normalized Lsm dwell time (**Fig. 4D**). In these plots, a lag phase is apparent for stable NTC accumulation for events in which sampling was observed as well as for those in which sampling was not detected. The NTC interacts transiently with pre-B/B complex spliceosomes containing Lsm2-8, but its stable integration can only take place after activation commences.

### Spliceosomes Containing Lsm8 and NTC are Short-Lived

We then analyzed the kinetic features of spliceosomes containing Lsm8, Cef1, and Syf1 (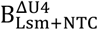 complexes) by identifying the arrival and departure times for each colocalized Lsm8-SNAP and Cef1/Syf1-DHFR event. The lifetimes of 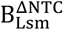 complexes were determined by subtracting the arrival times of Lsm8 signals from the arrival times of NTC signals 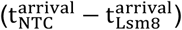, and the lifetimes of 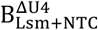 complexes were determined by subtracting the arrival times of NTC signals from the release times of Lsm8 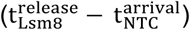 (**Fig. S9A**). Finally, the remaining lifetimes of complexes containing the NTC following Lsm8 release (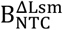 and later splicing complexes, L.C.) were found by subtracting the release time of Lsm8 from the release time of NTC 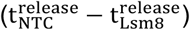. The distributions of dwell times were then fit as described above for the B_Lsm+U4_ and 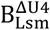 complexes.

In all cases, the distributions were best estimated using equations with a single exponential term as compared to more complex models using a loglikelihood ratio test (**Figs. S9B-D, Table S1**). The shortest-lived complex was 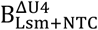 at 21.1±2.7 s. This is three-fold shorter than the lifetime of the 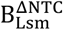 complex (68.6±8.9 s) and represents only about 5% of the average total lifetime of the NTC on the spliceosome. We also tested whether the individual lifetimes of any of the 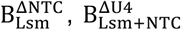 and 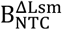 and subsequent complexes correlated with one another (**Figs. S6B-D**). As with U4 release, we could not identify any trends when the associated complex lifetimes were plotted against one another.

We previously determined that loss of U4 snRNP proteins during activation represented an irreversible step in splicing (19). Prp3/Prp4 almost never rejoined the pre-mRNA after their release. Instead, activated spliceosomes would either splice or disassemble to allow a different tri-snRNP molecule to associate and initiate a new activation process. In experiments with fluorescently labeled Lsm8 and Cef1/Syf1, Lsm8-SNAP rarely reappeared after its release while the NTC remained bound (4 out of 83 events). Consequently, U4 snRNP and Lsm2-8 release appear to happen sequentially and nearly irreversibly during splicing.

### Lsm2-8 Forms a Complex with U2/U6 helix II RNAs

Since Lsm2-8 is not released until after the NTC joins, we speculated that this could indicate an important function for Lsm2-8 during activation that is then taken over by the NTC. One possible function could be to maintain U2/U6 helix II structural integrity since it consists of only ten base pairs and has a calculated melting temperature of near 35°C in 1 M Na^+^ (30). In activated spliceosomes, this helix is surrounded by the NTC proteins Syf1 and Syf3 and directly bound by Syf2 (31, 32). These interactions may help stabilize helix II during the catalytic steps of splicing. In contrast, helix II is not known to be bound by proteins in B complex spliceosomes and is located adjacent to Lsm2-8 (**Fig. 5A**).

**Figure 5.**
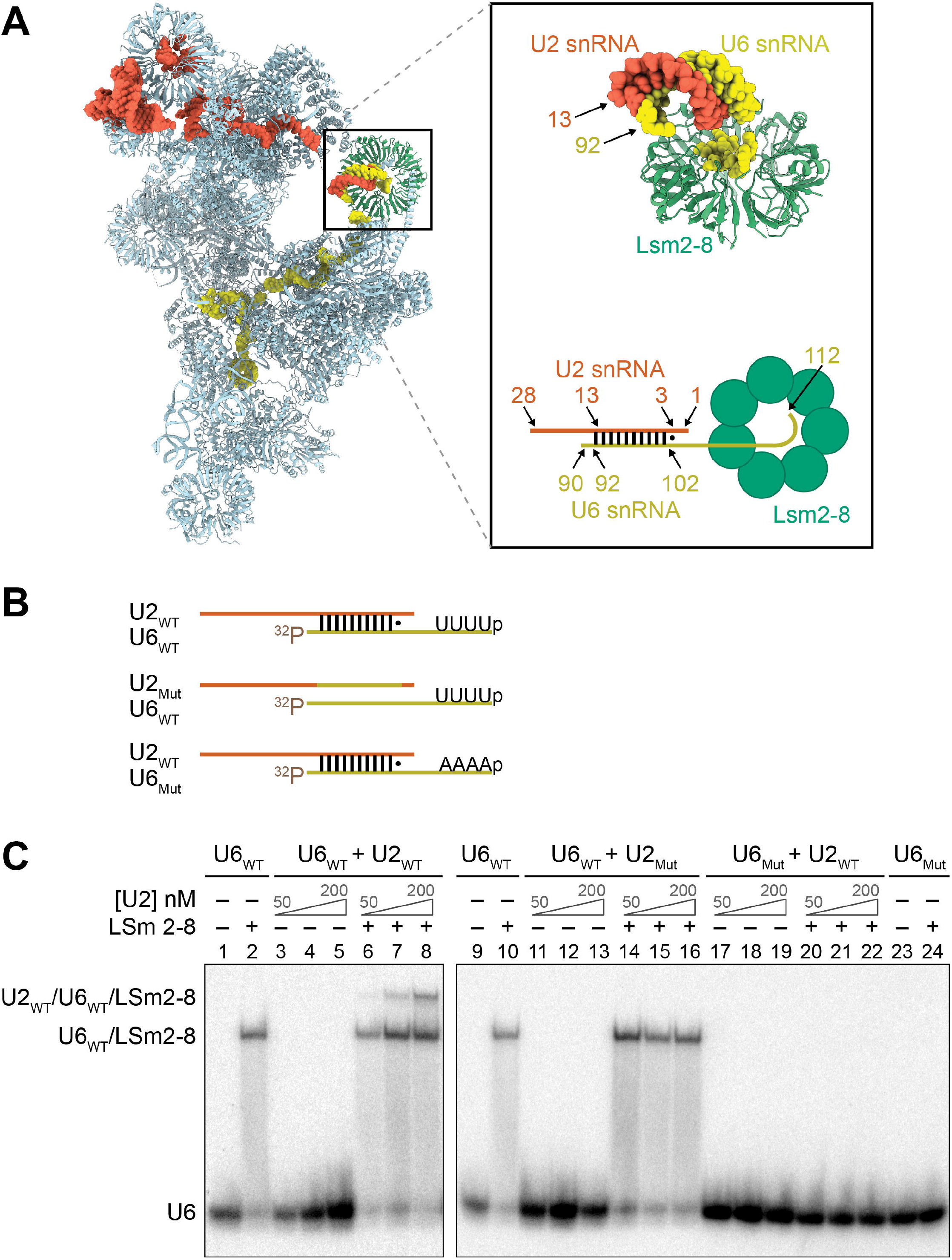
Formation of a Lsm2-8/U2/U6 ternary complex *in vitro*. (**A**) Cryo-EM structure (left) showing the location of Lsm2-8 (green), U2 snRNA (red) and U6 snRNA (yellow) in the yeast B complex spliceosome (PDB: 5NRL). Inset shows a close-up view of U2/U6 helix II located adjacent to Lsm2-8. The numbers of the terminal pairing nucleotides for U2 and U6 are indicated. Below the close-up view is a schematic of the structure with nucleotides that form helix II indicated. The U2_WT_ oligo used in EMSA assays represents nulceotides 1-28 of the U2 snRNA, and the U6_WT_ oligo represents nucleotides 90-112 of the U6 snRNA. (**B**) Cartoon illustating the U2 and U6 oligo pairs used for the *in vitro* binding assay (**Table S3**). All U6 oligos contain a 5’-[^32^P] label and a 3’-end phosphate for mimicking the processed 3’-end of yeast U6. The U6_Mut_ oligo does not contain a Lsm-binding site, and the U2_MUT_ oligo can not base pair to U6 to form helix II. (**C**) EMSA analysis of Lsm2-8/U2/U6 complex formation. Phosphor image of a native PAGE gel showing the presence of a U2-dependent supershifted complex (lanes 6-8). The supershifted complex does not appear if the U2 oligo cannot base pair to U6 (lanes 15-17) or if the U6 oligo lacks a Lsm-binding site (lanes 21-23). In all cases, the final concentrations of [^32^P]-labeled U6 oligos and Lsm2-8 were kept constant at 1nM and 250nM, respectively, while the U2 oligo concentration varied 50-200 nM if present.

We tested if Lsm2-8 could stabilize U2/U6 helix II *in vitro* with electrophoretic mobility shift assays (EMSAs) using purified recombinant Lsm2-8 and RNA oligo mimics of the 5’ and 3’ ends of the U2 and U6 snRNAs, respectively (**Fig. 5B**). As expected, Lsm2-8 tightly bound the U6_WT_ oligo (**Fig. 5C**, lane 2). When the U2 oligo was added, we observed a supershift dependent on the U2 oligo concentration (lanes 6-8). No supershift was observed if the U2 oligos could not base pair to U6 (lanes 15-17) or if the U6 oligo was missing a Lsm2-8 binding site (lanes 21-23). Together, the data show that Lsm2-8 can form a ternary complex with the U2 and U6 RNAs dependent on base pairing and Lsm2-8 binding to U6. Lsm2-8 may function in spliceosomes to maintain U2/U6 helix II until it can be stabilized by NTC proteins, possibly by handover from Lsm2-8 to the NTC in the 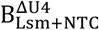 complex.

## DISCUSSION

### A Kinetic Scheme for Spliceosome Activation

By integrating our experimental data, we can define a kinetic scheme for yeast spliceosome activation *in vitro* (**Fig. 6**). In the scheme, the tri-snRNP first joins the spliceosome A complex to form the pre-B and subsequent B complex. The average combined lifetime of these complexes is ∼42 s. Multiple rearrangements likely occur during this time including release of the U1 snRNP, which has not yet been kinetically characterized. Following the loss of U4 snRNP, a 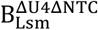 complex likely forms, which we define as containing the Lsm2-8 ring but lacking the U4 snRNP and NTC. To our knowledge, this complex has not previously been reported for either yeast or human spliceosomes. While we did not directly observe this intermediate in our experiments, we can infer its existence from previous data showing that the NTC is primarily recruited after U4 snRNP release (19) and experiments reported here showing that Lsm2-8 release occurs after NTC recruitment (**Fig. 3**). The lifetime of the 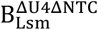 intermediate can be estimated by either subtracting the lifetime of 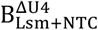 from the lifetime of 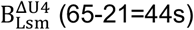 or by subtracting the lifetime of B_Lsm+U4_ from the lifetime of 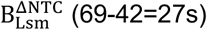 (**Table S1**). This yields a range of 27-43s for the lifetime of 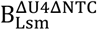.

**Figure 6.**
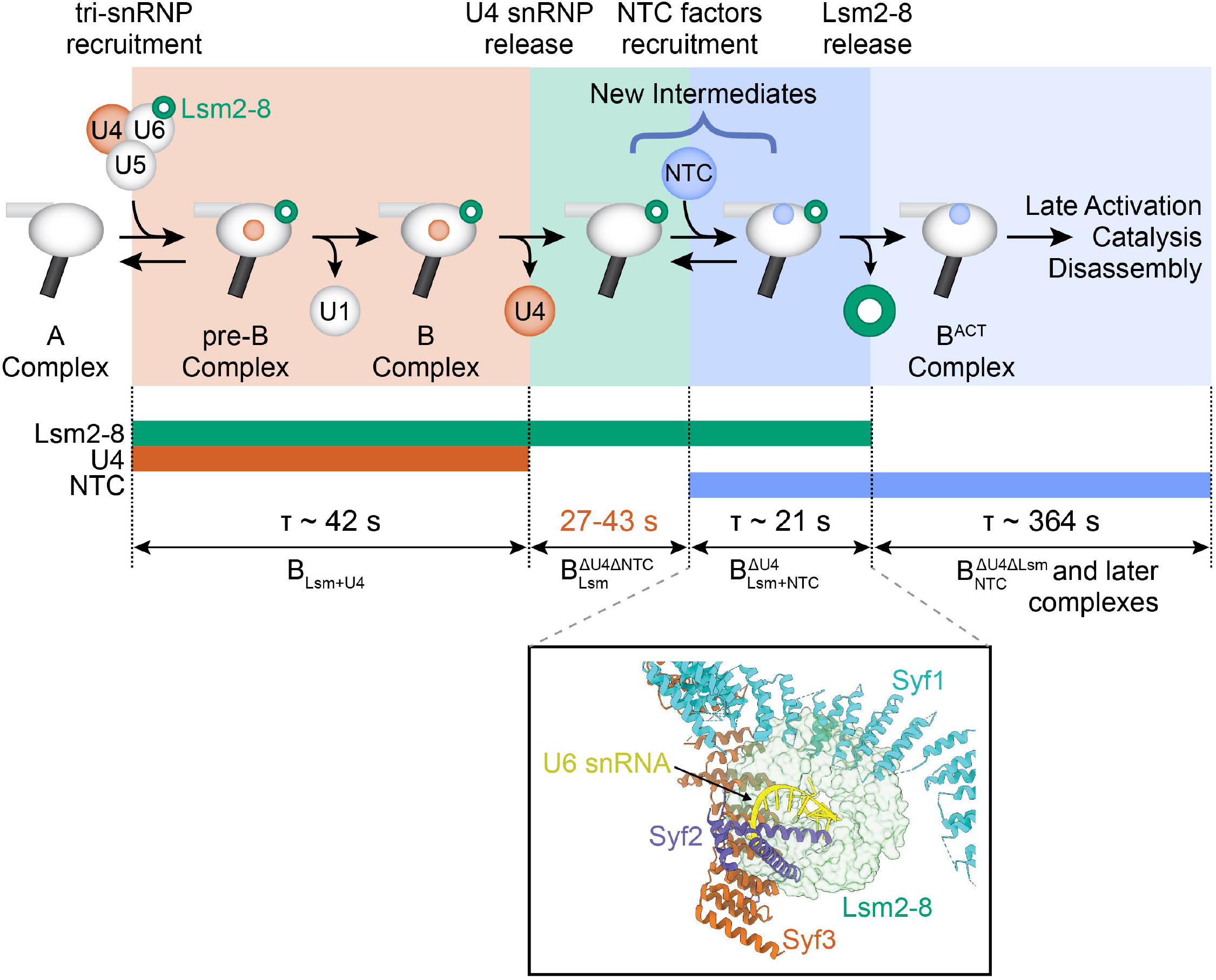
Transient Intermediates Formed During Spliceosome Activation. In this kinetic scheme, U4 snRNP release precedes NTC recruitment, which in turn precedes Lsm2-8 release. This would involve formation of at least two activation intermediates, the 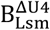 and 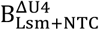 complexes, which have not been previously biochemically or structurally characterized. Single molecule data allows determination of the characteristic lifetimes (ρ) of these complexes. The 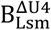 complex lifetime (red) was not directly measured in our experiments but can be inferred as described in the text (19). Not pictured in this scheme are NTC sampling events that begin soon after tri-snRNP recruitment. (*Inset*) The structure of the 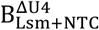 complex represents a unique conformation of the spliceosome since the Lsm2-8 ring and the NTC proteins Syf1 and Syf3 are mutually exclusive with one another when the structures of yeast B and B^ACT^ complexes are superimposed (PDBs: 5NRL and 5GM6).

Subsequently, the NTC is recruited to 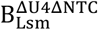 to form the 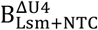 spliceosome, which persists for only ∼21 s before the release of Lsm2-8 ring. Like the preceding complex, 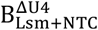 has not previously been observed for yeast spliceosomes but shares some similarities with the human pre-B^ACT1^. Following this step, the NTC typically remains associated for ∼364 s. This remaining NTC lifetime is likely a composite of subsequent steps in splicing and spliceosome discard. Our kinetic scheme also supports the hypothesis that the human pre-B^ACT1^ complex observed in the presence of splicing inhibitors (18) is a competent activation intermediate. We were able to detect spliceosomes compositionally similar to human pre-B^ACT1^ being formed and transformed at rates rapid enough to be consistent with *in vitro* splicing. Overall kinetic efficiency of the activation process may be achieved, in part, by the presence of at least two irreversible steps: ordered release of the U4 snRNP and Lsm2-8.

### Activation Intermediates Suggest New Spliceosome Conformations

It is likely that the 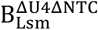 and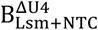 spliceosomes have unique conformations not yet observed in cryo-EM spliceosome structures. No structural information yet exists for spliceosomes lacking both U4 snRNP and NTC; however, biochemical evidence for such a complex has been obtained for the human splicing machinery in the presence of inhibitors (33). Lsm2-8 and NTC proteins must be uniquely positioned in the 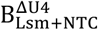 spliceosome since these factors are mutually exclusive when the structures of yeast B and B^ACT^ complexes are superimposed (**Fig. 6B**). The structure of 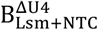 is also distinct from that of human pre-B^ACT1^ complex since our data indicate that Syf1 is present in B_Lsm+NTC_ but this protein is missing from pre-B^ACT1^. Indeed, it was noted that human Syf1 cannot bind pre-B^ACT1^ complexes due to a mutually exclusive binding site with Lsm2-8 (18). Whether or not a transient human spliceosome containing both Syf1 and Lsm2-8 (analogous to yeast 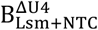) is formed during the pre-B^ACT1^ to pre-B^ACT2^ transition is not yet known.

### NTC Samples Spliceosomes during Activation

Watching NTC recruitment in real time, we were able to observe transient NTC binding events to B complex spliceosomes that precede stable NTC integration. Since these transient events are more likely to occur early in activation and stable events more likely to occur later, it is consistent with transition of the spliceosome from a low to high affinity NTC-binding state. We have not identified the transition required for high affinity binding, but it could involve structural remodeling permitted by U4 snRNP release to allow the spliceosome to accommodate both the NTC and Lsm2-8.

The sampling behavior itself may allow the NTC to surveil spliceosomes for efficient and correct progress through activation. Rapid and reversible NTC binding may prevent NTC complexes from being sequestered by malfunctioning spliceosomes such as those which fail to release U4. Since the NTC is also involved in nuclear processes other than splicing (34-36), these features of NTC binding dynamics might be important for preventing depletion of the available NTC pool by limiting formation of stable but unproductive complexes.

### Divergent Pathways for NTC Binding in Yeast and Human Spliceosomes

Recently, it has been proposed that the human NTC is composed of discrete subunits that associate with the spliceosome during activation in a stepwise mechanism (18). Initially, the hNTC subunit containing CDC5L (yeast Cef1) and five other factors associates to form the pre-B^ACT1^ complex. In a second step, the intron binding complex (IBC) that contains SYF1 along with the hNTC-related (hNTR) proteins and other factors associate to form the pre-B^ACT2^ complex. In contrast, biochemical and mass spectrometry data support the existence of a single yeast NTC complex containing Cef1, Syf1, Prp19 along with IBC and NTR components (37, 38).

Our single molecule data are also consistent with the existence of a NTC complex containing Cef1 and Syf1. While it was previously speculated that the smaller size of the hNTC relative to its yeast counterpart permits the simultaneous presence of hNTC with Lsm2-8 during activation (18), our results show that this is still possible even with the larger yeast NTC, likely due to structural rearrangements in the NTC or Lsm2-8 binding region. Thus, stable NTC association with the spliceosome in yeast appears to involve both fewer steps and individual subcomplexes than in humans. This may contribute to greater efficiency of yeast splicing but at the expense of fewer potential points at which splicing can be regulated during activation.

### A Function for Lsm2-8 in Maintaining U2/U6 Helix II during Activation

One activity that could occur within the 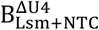 complex is transfer of the relatively unstable U2/U6 helix II to the NTC. Since Lsm2-8 is located adjacent to U2/U6 helix II, it could facilitate formation of this RNA duplex and/or help to stabilize it once formed. Avoiding release of Lsm2-8 in the absence of NTC binding may therefore be important for maintaining helix II integrity during activation and preventing spurious unwinding which could lead to spliceosome disassembly. Consistent with this hypothesis, we observed that purified Lsm2-8 can form a ternary complex with U2/U6 helix II RNAs dependent on base pairing (**Fig. 5**). How important this function is for the yeast splicing machinery is unclear since neither helix II nor the helix II-interacting NTC protein Syf2 are essential for yeast viability (39, 40). Lsm2-8 stabilization of helix II during activation may only become essential under certain conditions in which the nascent spliceosome catalytic center is destabilized. Indeed, helix II becomes essential when the spliceosome catalytic center is mutated (39). Interestingly, Lsm2-8 also facilitates annealing of the U4 and U6 snRNAs by Prp24 (41). In this case, it was proposed to occur through Lsm2-8-facilitated folding of the intramolecular U6 telestem RNA duplex. The 3’ half of the telestem ultimately pairs with U2 to form U2/U6 helix II. It is possible that similar properties of Lsm2-8 contribute to either U6 telestem or U2/U6 helix II stabilization, both of which involve duplex formation to the same region of U6 upstream of the Lsm2-8 binding site.

Formation of a complex between two complementary RNAs (U2 and U6) and Lsm2-8 is also intriguing due to similarities between eukaryotic Lsm proteins and bacterial Hfq. Even though Hfq proteins assemble into homohexamers and not heteroheptamers like Lsm2-8, both Hfq and Lsm proteins share a common fold (10). Hfq has been extensively studied for its ability to anneal bacterial small RNAs to mRNAs to regulate their expression (42). Our data suggest that in addition to a common fold, interaction with unstable RNA duplexes has also been evolutionarily conserved between the Lsm and Hfq proteins. The spliceosome may use these interactions to help form and/or stabilize U2/U6 helix II through assembly and the conformational and compositional changes occurring during activation. Whether or not Lsm1-7 or Sm complexes also possess similar duplex stabilizing activities is not yet known but could yield additional functional parallels to Hfq.

## SUPPLEMENTARY DATA

Supplementary Information accompanies this article.

## ACKNOWLEDGEMENTS

We thank Karli Lipinski and David White for critical reading of the manuscript and members of the Hoskins, Brow, and Butcher laboratories for helpful discussions.

## AUTHOR CONTRIBUTIONS

XF, MLR, and AAH conceived the project. XF, HK, and MLR carried out experiments. ECM and SEB provided purified Lsm proteins. XF, HK, MLR, and AAH analyzed data. XF and AAH wrote the manuscript with input from all authors.

## MATERIAL AND METHODS

### RNA Oligonucleotides

RNA oligos (**Table S3**) were purchased from Integrated DNA Technologies. Stocks (100 µM) were prepared by resuspending the oligos in nuclease-free water (Ambion). Initial stock concentrations were calculated from their absorbance values and the predicted extinction coefficients at 260nm using a NanoDrop.

### Yeast Strains

Yeast strains (**Table S4**) were derived from the protease-deficient strain BJ2168 and splicing factors were c-terminally tagged by integrating fast SNAP or DHFR tags at the appropriate genomic locations by homologous recombination as previously described (19, 20).

### Recombinant Lsm2-8 Expression and Purification

Lsm2-8 proteins were expressed in *E. coli* from a single plasmid and the heteroheptamer was purified as previously described (16).

### 5’-end Labeling of RNAs

Oligos (10 pmols for U6) were 5’-end labeled with [γ-^32^P]-ATP (PerkinElmer), using T4 PNK and 10X reaction buffer A (Thermo Fisher Scientific) at 37°C for 1 h, then heat inactivated at 75°C for 10 min. The labelled oligos were then gel purified on a 19-20% PAGE denaturing gel (AccuGel 19:1, National Diagnostics), followed by extraction, ethanol precipitation, resuspension in buffer and quantification by liquid scintillation counting.

### Electrophoretic Mobility Shift Assays (EMSAs)

Complexes between U2 and U6 RNAs and the Lsm2-8 protein were formed in a volume of 10 μL by mixing U2 and U6 oligos in RNA dilution buffer (100mM KCl, 20% w/v sucrose, 20mM HEPES (pH 7), 1mM EDTA (pH 8), 1mM TCEP·HCl, 0.01% v/v Triton X-100, 0.2 mg/mL yeast tRNA [Ambion, Catalog No. AM7119], 0.2 mg/mL sodium heparin [Sigma Aldrich, Catalog No. H4784]) together with purified Lsm2-8 in protein dilution buffer (100mM KCl, 20% w/v sucrose, 20mM HEPES (pH 7), 1mM EDTA (pH 8), 1mM TCEP·HCl, 0.01% v/v Triton X-100, 0.2mg/mL BSA [Pierce, Catalog No. 23209]) in equal volume. Final concentrations for U6, U2 oligos and purified Lsm2-8 are described in the corresponding figures. All reactions were incubated at room temperature (22-24°C) for 30 min prior to gel electrophoresis.

After 30min, the reactions were placed on ice and then loaded onto a native 6% polyacrylamide gel (AccuGel 29:1, National Diagnostics), which had been pre-run with 1x TBE buffer at 300 V for 30 min at 4°C. Electrophoresis was then carried out at 300 V for 2-4 h. For detection of radioactive bands, the gels were dried and then exposed to a phosphor screen. Phosphor images of the screens were taken in the phosphorescence mode of a Typhoon scanner (Cytiva). Results were analyzed with ImageQuantTL (Cytiva) software.

### Preparation of Yeast WCE

Yeast WCE for splicing was prepared according to published protocols using the ball method (43). SNAP-tagged proteins were fluorophore labeled as previously described (44). Briefly, SNAP-Surface® 549 dye (S9112S, New England BioLabs; abbreviated elsewhere as SNAP-DY-549) in DMSO was added to 1.2 mL yWCE to a final concentration of 1 μM. The reaction tube was rotated in the dark for 30 min at room temperature. The reaction was then loaded onto a pre-equilibrated G-25 Sephadex column (Kontes Flex Column) in SEC buffer (25 mM HEPES-KOH pH 7.9, 50 mM KCl, 1 mM DTT, 10% v/v glycerol) at 4°C to remove excess dye. A Peristaltic Pump P-1 (Cytiva) was used for pre-equilibrating the column and eluting the labeled extract at a flow rate of ∼0.25 mL/min with the SEC buffer. Fluorophore labeling of the proteins was confirmed by SDS-PAGE and fluorescence using a Typhoon FLA 9000 scanner (Cytiva) at 532nm. Results were analyzed with ImageQuantTL (Cytiva) software.

### In Vitro Splicing Assays

[α-^32^P] UTP radio-labeled (PerkinElmer) and m^7^G(5’)ppp(5’)G capped (New England Biolabs) RP51A pre-mRNA substrates were made by *in vitro* transcription of a linear DNA template with T7 RNA polymerase (Agilent or purified in the laboratory). The DNA template was produced from a PCR reaction of pBS117 plasmid (45) using Taq DNA polymerase (M7122, Promega), followed by gel purification of the products with a Wizard SV Gel and PCR Clean-Up System kit (Promega). Transcription products were separated on a 6% denaturing polyacrylamide gel (AccuGel 19:1, National Diagnostics), followed by ethanol precipitation of the extracted RP51A transcripts and quantitation of the resuspended transcripts in nuclease-free water (Ambion, Fischer Scientific) with a liquid scintillation counter (LSC, tri-carb 2900TR, Packard).

A typical *in vitro* splicing reaction included 40% v/v WCE and 0.2 nM radio-labeled RP51A substrate in a splicing buffer [final concentrations: 100mM KPi pH 7.3, 3% w/v PEG-8000, 1mM DTT, 2.5mM MgCl_2_, 0.2U/μL RNasin Plus (Promega)]. The reaction was incubated at room temperature for 45 min. The reaction was quenched in a splicing dilution buffer (100mM Tris base pH 7.5, 10mM EDTA pH 8.0, 1% w/v SDS, 150mM NaCl, 300 mM NaOAc pH 5.3). RNAs from the reaction were extracted using phenol-chloroform, ethanol precipitated, resuspended in deionized formamide, and separated on a 12% denaturing polyacrylamide gel. Gels were then dried and exposed to a phosphor screen overnight. The screen was imaged with a Typhoon FLA 9000 scanner (Cytiva). Results were analyzed with ImageQuantTL (Cytiva) software. The intensities (*I*) for RP51A pre-mRNAs and splicing products were determined by integrating the signals within same-sized rectangles around the band. The background corrected intensities for bands were then used for calculating splicing efficiencies, with Equations 1 and 2.

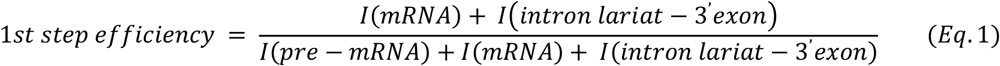

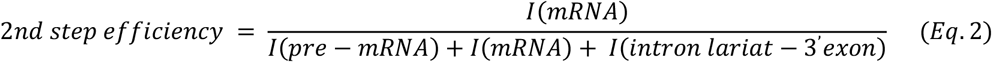

### Preparation of Fluorescently-Labeled RP51A pre-mRNAs

Unlabeled biotin handles (**Table S3**, Oligo 8) from IDT were dissolved in 0.091 M Na_2_B_4_O_7_ buffer (NaB buffer, pH 8.50, adjusted with concentrated HCl) at a concentration of 1 mM. Alexa Fluor 488 NHS ester (A20000, Thermo Fisher Scientific) was prepared by dissolving to a concentration of 1 mg in 60 µL of anhydrous DMSO. For fluorophore labeling of the biotin handle, 15 µL of the fluorophore was mixed with 49 µL of the biotin handle and 36 µL NaB buffer. The reaction was kept in the dark and rotated overnight at room temperature. The labeling reaction mixture was then applied to a G-25 MicroSpin column (45-001-397, Fisher Scientific) to remove most of the unreacted dye. The eluate from the spin column was then loaded onto a 19% denaturing polyacrylamide gel (AccuGel 19:1, National Diagnostics). Fluorescent bands containing the oligo were excised from the gel, the oligo eluted from gel, and then ethanol precipitated. The products were resuspended in water and concentrations determined by UV-Vis.

Capped RP51A transcripts were made by *in vitro* transcription using T7 RNA polymerase and the biotin handle attached by splinted RNA ligation. The fluorescently labeled oligo (56 pmol) was 5’-phosphorylated using T4 PNK (5U, M0201S, New England Biolabs) according to the manufacturer’s directions with the addition of 2 mM ATP. The PNK enzyme was then heat inactivated and DNA bridge oligos (50 pmol; **Table S3**, Oligo 9) and radio-labeled RP51A transcript (14 pmol) was added and then allowed to anneal by incubating at 90°C for 3 min, followed by cooling using a gradient of 0.1°C/s over 10 min. T4 RNA Ligase 2 (20U; M0239L, New England Biolabs) was then added to ligate the transcript to the biotinylated oligo. Ligation products were separated on a 5% denaturing polyacrylamide gel (AccuGel 19:1, National Diagnostics), followed by ethanol precipitation of the extracted products from the gel. Precipitated RNAs were resuspended in nuclease-free water (Ambion, Fischer Scientific) and quantified with a liquid scintillation counter (LSC, tri-carb 2900TR, Packard).

### CoSMoS Assays

Microscope slides (100490-396, VWR) and cover glasses (12-553-455, Fischer) were cleaned by sonication while immersing in the following solutions/solvents in the order listed: f 2% v/v micro-90, 100% ethanol, 1M KOH, and MiliQ water. Each sonication step took 1 h, followed by rinsing with MiliQ water between steps. After drying with high purity nitrogen (NI UHP300, Airgas), the slides were aminosilanized with VECTABOND (NC9280699, Fisher Scientific). Reaction chambers were created by drawing multiple parallel straight lines on the slides with vacuum grease, followed by covering with the cover glasses to create individual lanes for reaction chambers. Typically, four lanes can be made on a single slide. Lastly, the slides were passivated by addition of a mixture of mPEG-SVA (MPEG-SVA-5K, Laysan Bio) and mPEG-biotin-SVA (BIO-PEG-SVA-5K, Laysan Bio) at a ratio of 1:100 w/w in 100mM NaHCO_3_ (pH 8) buffer and incubating overnight.

Prior to each CoSMoS assay, the mPEG mixture in the lane (∼20 μL) was washed off with 1x PBS (200 μL) twice. Then, streptavidin (50 μL, 0.2 mg/mL, SA10, Kelowna International Scientific) was flowed in the lane, allowing it to bind with the biotin on the slide surface. After washing the lane off with 200 μL 1x PBS, 50 μL 50-150 pM fluorescently labeled and biotinylated RNA was added and allowed to bind with the streptavidin. The slide was washed again with PBS and the density of RNA was determined by exciting the fluorophore with the 488 nm laser. Finally, splicing assay buffer containing 40% v/v WCE, 20 nM Cy5-TMP (20), oxygen scavenger (5 mM PCA and 0.96 U/mL PCD), 1 mM Trolox, and triplet quenchers (0.5 mM propyl gallate, 1 mM cyclooctatetraene, 1 mM 4-nitrobenzyl alcohol) was added (20). The triplet quenchers were added as a mixture made as a 100x stock in DMSO.

### CoSMoS Data Acquisition

A custom-built objective-type TIRF microscope (23, 46) together with Glimpse software (written in LabVIEW programming language, https://github.com/gelles-brandeis/Glimpse) was used for collecting single molecule movies. Briefly, a 60X 1.45(NA) PlanApo objective (Olympus) was used for the TIRF excitation and collecting the emission light. Different wavelengths of incoming and exit excitation light were directed to and away from the objective by two separate micromirrors. The emission light was further filtered and separated by a dichroic mirror at the cutoff wavelength of 635 nm. The spectrally separated light was then focused onto two different areas (FOV: field of view) of the same EMCCD camera. Four different lasers were used in this study, including 488 nm (blue), 532 nm (green), 633 nm (red) and 785 nm (infrared), the powers of which were set in the ranges of 1.2 – 1.5 mW, 500 – 600 μW, 400-440 μW, and 2.5 mW, respectively. Cycles of time-lapse imaging were used according to the following excitation scheme with each frame lasting 1s: in each cycle, the 785 nm was first used to illuminate the sample and correct sample positioning using an auto-focus system. Then, the 488 nm blue laser was turned on to collect two consecutive frames to image the immobilized RNAs. The 532 and 633 nm lasers were then turned on to simultaneously collect 14 frames with a 3 s delay between adjacent frames. The total cycle time was ∼1 min and this cycle was repeated 50x to collect videos lasting for ∼50 min. To avoid photo bleaching of DY-549 and Cy5 fluorophores by the 488 nm laser, the path of the laser was physically blocked to prevent the beam from reaching the sample after collection of 10 frames of blue images.

### CoSMoS Data Analysis

Single molecule data were analyzed by the same method as described previously (20, 28) with slight modifications, using a custom program imscroll (https://github.com/gelles-brandeis/CoSMoS_Analysis) written in MATLAB (The Mathworks). A general analysis procedure includes, 1) creating a mapping file for correlating the locations in the <635 nm FOV to those in the >635 nm FOV, using a fluorescent beads (T10711, ThermoFisher); 2) creating a driftlist file for correcting the drift of the fluorescent spots over the recording; 3) creating a AOI (area of interest; typically either 3×3 or 5×5 pixels) file listing the positions of the immobilized pre-mRNA molecules; 4) combining the AOI, mapping, and driftlist files to generate drift-corrected AOIs corresponding to the pre-mRNA locations in the fields of view used for DY-549 and Cy5-TMP imaging; and 5) integrating the measured intensities in the AOIs in those fields of view over the experimental time course (47). Binding events were identified as signals centered within the AOI that appeared with intensities greater than 3.6x the standard deviation above the mean of the background noise. Loss of signals were identified as points in time at which the signal fell below 1x standard deviation above the mean background.

Analysis of the measured dwell times and fits to kinetic equations were carried out using MATLAB and AGATHA software (https://github.com/hoskinslab/AGATHA) using maximum likelihood methods and fitting to equations containing one (Eq. 3), two (Eq. 4), or three (Eq. 5) exponential terms as described (28).

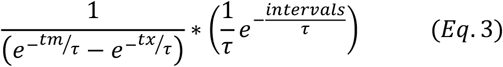

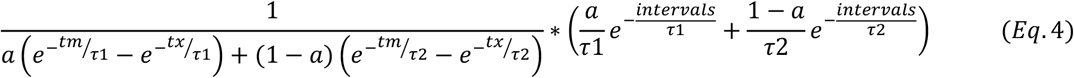

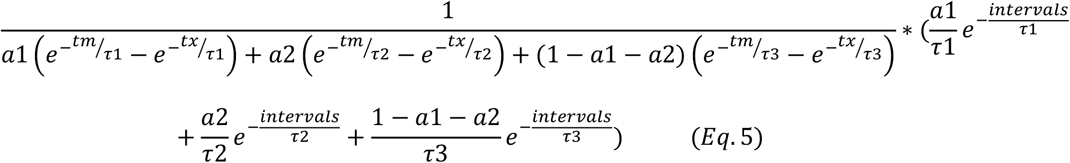

Bootstrapping was used to calculate standard errors for all fitted parameters. Fit parameters are included in **Table S1**, and models that best fit the distributions were identified by using the log-likelihood ratio test (28).

Histograms for the distribution of events were generated in MATLAB with empty bins removed. The error (standard deviation) for each bin were calculated using Eq. 6, with the assumption that the number of events within a bin follows a binomial distribution.

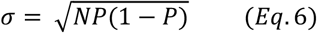

