## Supplemental Information for "Identification of Transient Intermediates During Spliceosome Activation by Single Molecule Fluorescence Microscopy"

**This PDF includes:**

9 Supplemental Figures and Figure Legends

4 Supplemental Tables

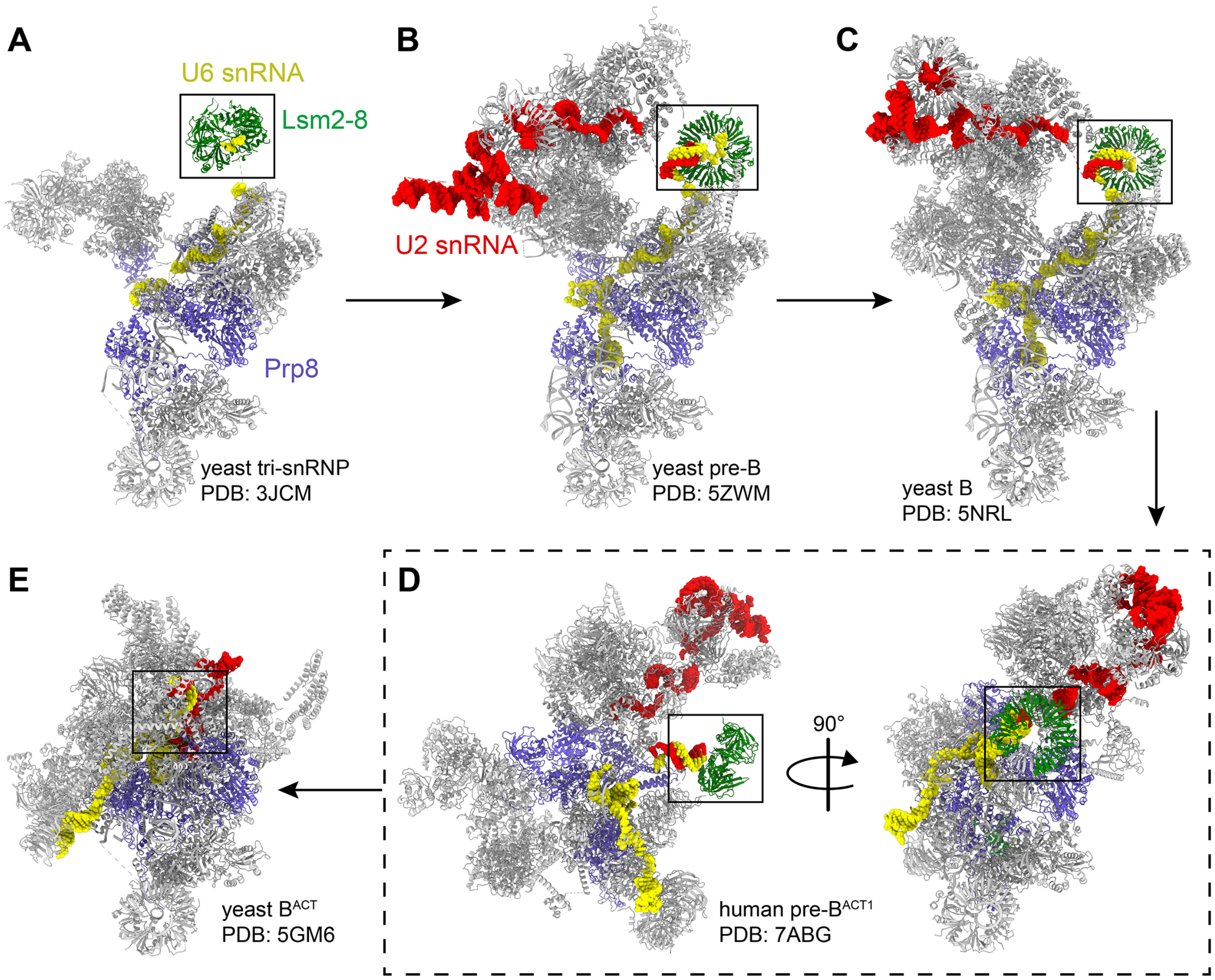


**Figure S1. Structural Scheme for Spliceosome Activation.** Conformational and compositional changes occurring during spliceosome activation beginning with the yeast U4/U6.U5 tri-snRNP (**A**) and ending with the yeast $B^{ACT}$ complex (**E**). The box in each structure is centered on the 3' end of the U6 snRNA while structures have been roughly aligned based on the position of the U5 snRNP Sm ring (grey ring at the base of each structure). The structures in panel (**D**) represent the human, not yeast, pre-$B^{ACT1}$ spliceosome since no intermediate structures have yet been obtained for the yeast B to $B^{ACT}$ transition.

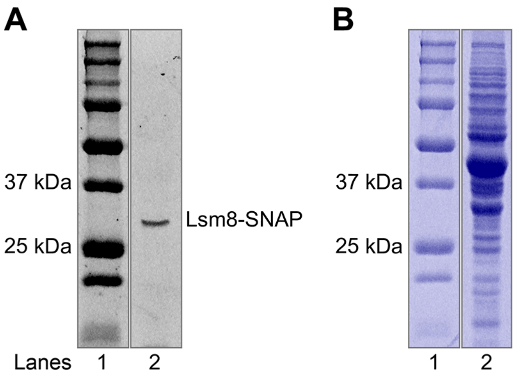


**Figure S2. Fluorophore Labeling of Lsm8-SNAP.** Lsm8-SNAP proteins can be specifically labeled in yeast WCE with SNAP-DY-549. Shown is fluorescence image (**A**) and Coomassie stain (**B**) of the same SDS-PAGE gel used to analyze Lsm8-SNAP-containing extract after fluorophore labeling. The gel images were cropped (black boxes) to remove intervening lanes not relevant to this figure.

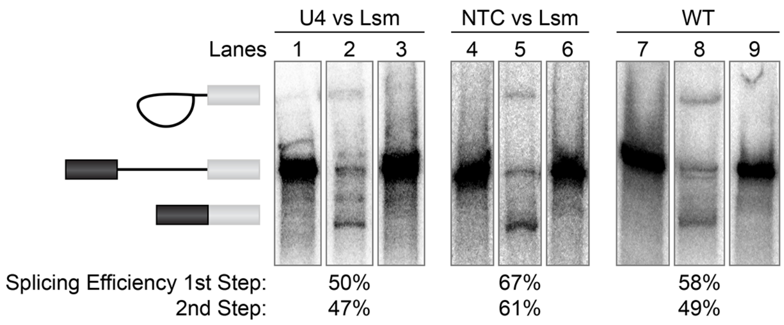


**Figure S3. Splicing Activity of Lsm8-Labeled WCE.** Representative splicing assays and efficiencies for WCE containing Lsm8-SNAP labeled proteins (lanes 1-6) or a WT control (lanes 7-9). In lanes 2, 5, and 8, ATP was added to a concentration of 2 mM to promote splicing. In lanes 3, 6, and 9 ATP was added to a concentration of 0.05 mM to permit spliceosome assembly but prevent activation and splicing. Lanes 1, 4, and 7, represent samples at t=0 min while the other lanes represent reactions quenched at 45 min. The gel images were cropped (vertical white stripes) and combined into a single figure to remove intervening lanes not relevant for this figure and for clarity.

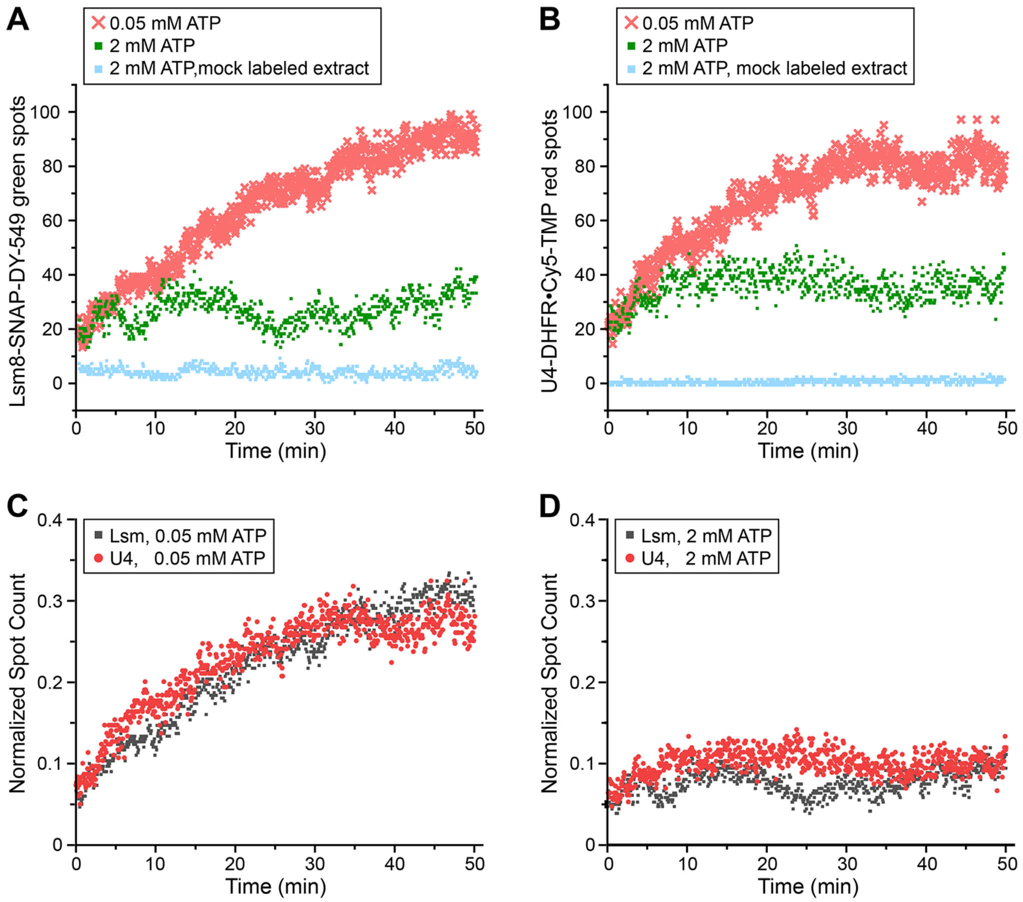


**Figure S4. ATP-Dependent Accumulation of Lsm8 and U4 Protein Fluorescent Spots.** Shown are spot accumulation trajectories for CoSMoS experiments containing green-labeled Lsm8 and red-labeled U4 proteins. In panels (**A** and **B**), spot accumulation trajectories are compared for the same experiment from both the green/Lsm8 (**A**) and red/U4 (**B**) channels under conditions that inhibit (0.05 mM ATP) or permit activation and splicing (2 mM ATP). As a control, the spot accumulation trajectory is shown for a WT extract labeled with SNAP-DY-549 and Cy5-TMP fluorophores but not containing any SNAP- or DHFR-tagged proteins (blue). In panels (**C** and **D**), normalized spot counts from the data shown in panels (**A** and **B**) were calculated by dividing the number of observed fluorescent spots by the number of surface-immobilized pre-mRNAs in the field-of-view. Data for Lsm and U4 were then overlaid at low (**C**) and high (**D**) ATP to highlight the similarities in trajectories for the splicing factors.

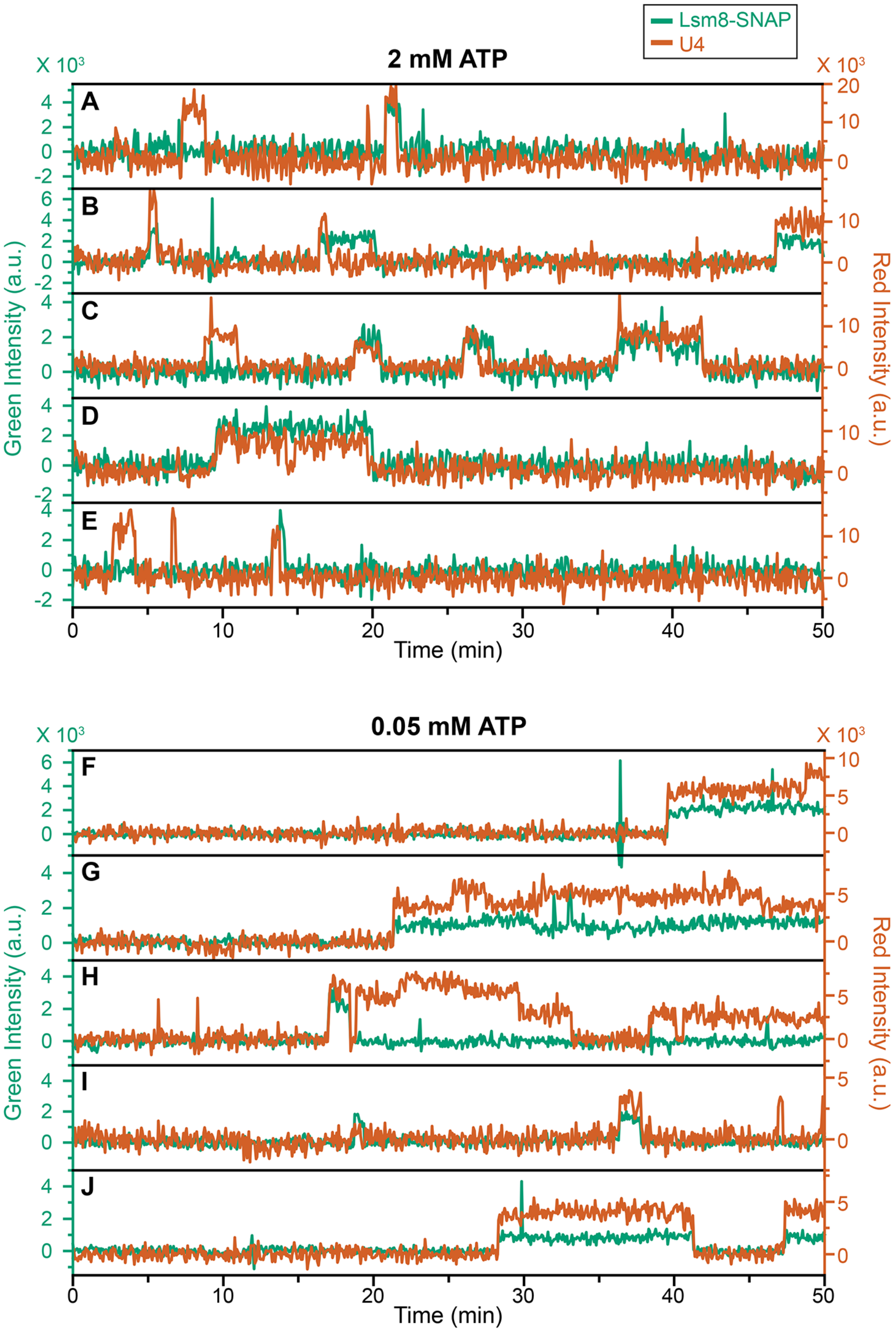


**Figure S5. Additional Sample Fluorescence Trajectories from 3-color CoSMoS Experiments Monitoring Lsm8-SNAP and U4-DHFR Proteins.** Shown are super-imposed fluorescence intensities for Lsm8-SNAP (green traces) and U4-DHFR proteins (red traces) at 2 mM ATP (**A-E**) and 0.05 mM ATP (**F-J**).

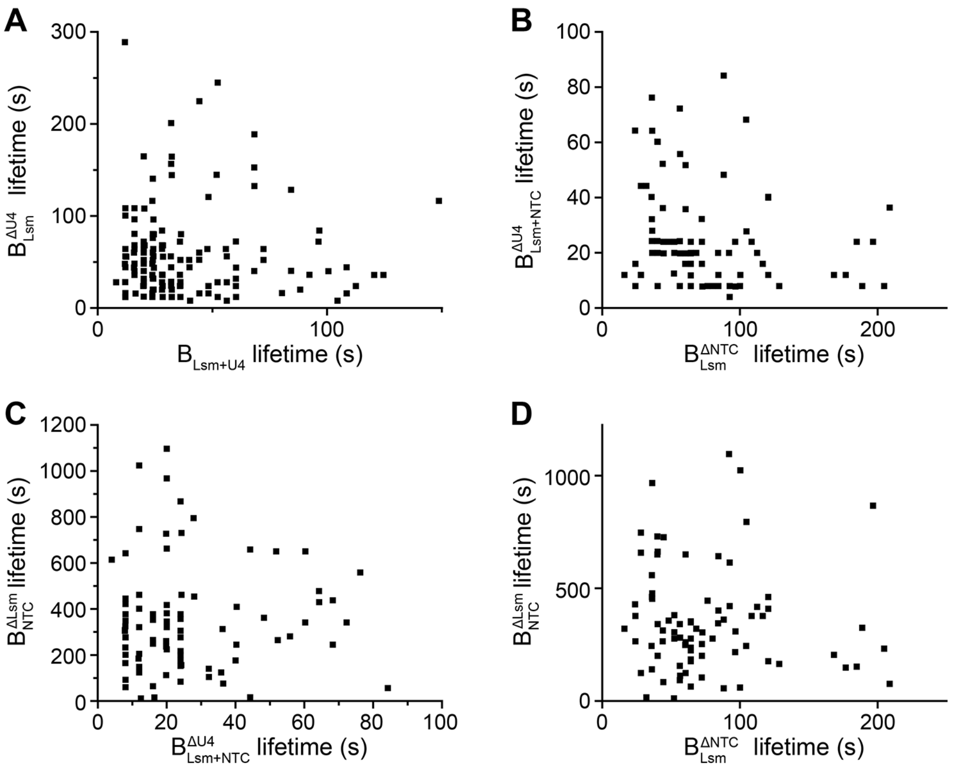


**Figure S6. Lifetimes of Spliceosome Complexes Exhibit Little Correlation with One Another.** The lifetimes for paired events for the indicated spliceosome complexes obtained from 3-color CoSMoS experiments were plotted against one another. (**A**) Correlation of *N*=144 pairs of lifetimes for the indicated complexes. For this dataset, a Pearson's correlation coefficient (r) of 0.17 was observed. (**B**) Correlation for *N*=84 pairs of lifetimes yielded r = -0.09. (**C**) Correlation for *N*=83 pairs of lifetimes yielded r = 0.02. For this data set one outlier was removed with a NTC lifetime of >2000 s. (**D**) Correlation for *N*=84 pairs of lifetimes yielded r = -0.15.

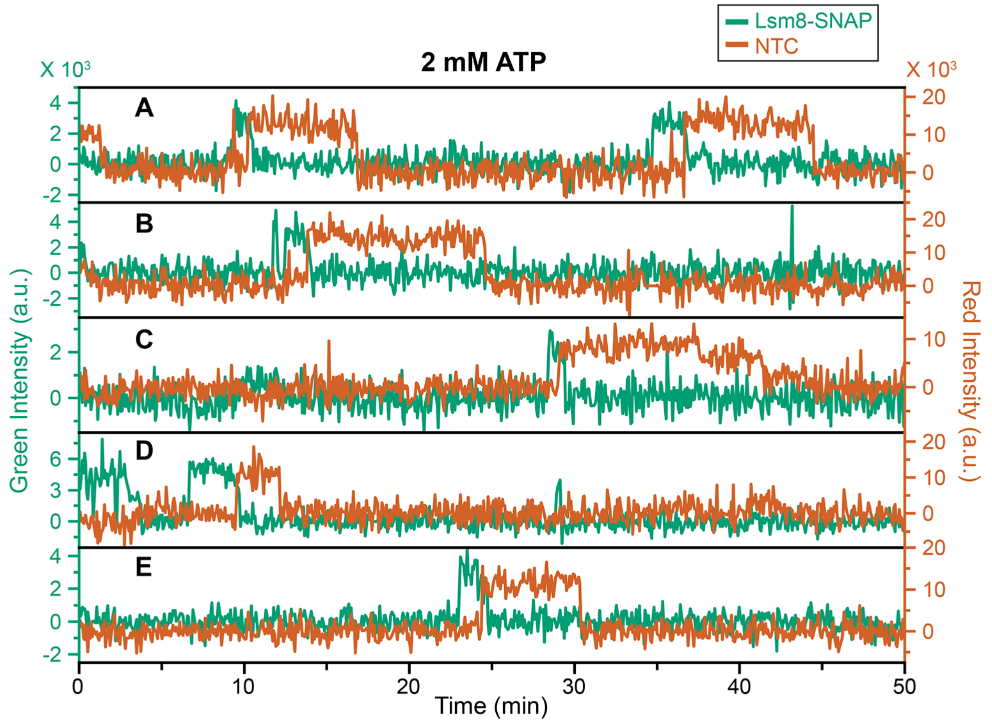


**Figure S7. Additional Sample Fluorescence Trajectories from 3-color CoSMoS Experiments Monitoring Lsm8-SNAP and NTC-DHFR Proteins.** Shown are super-imposed fluorescence intensities for Lsm8-SNAP (green traces) and NTC-DHFR proteins (red traces) at 2 mM ATP.

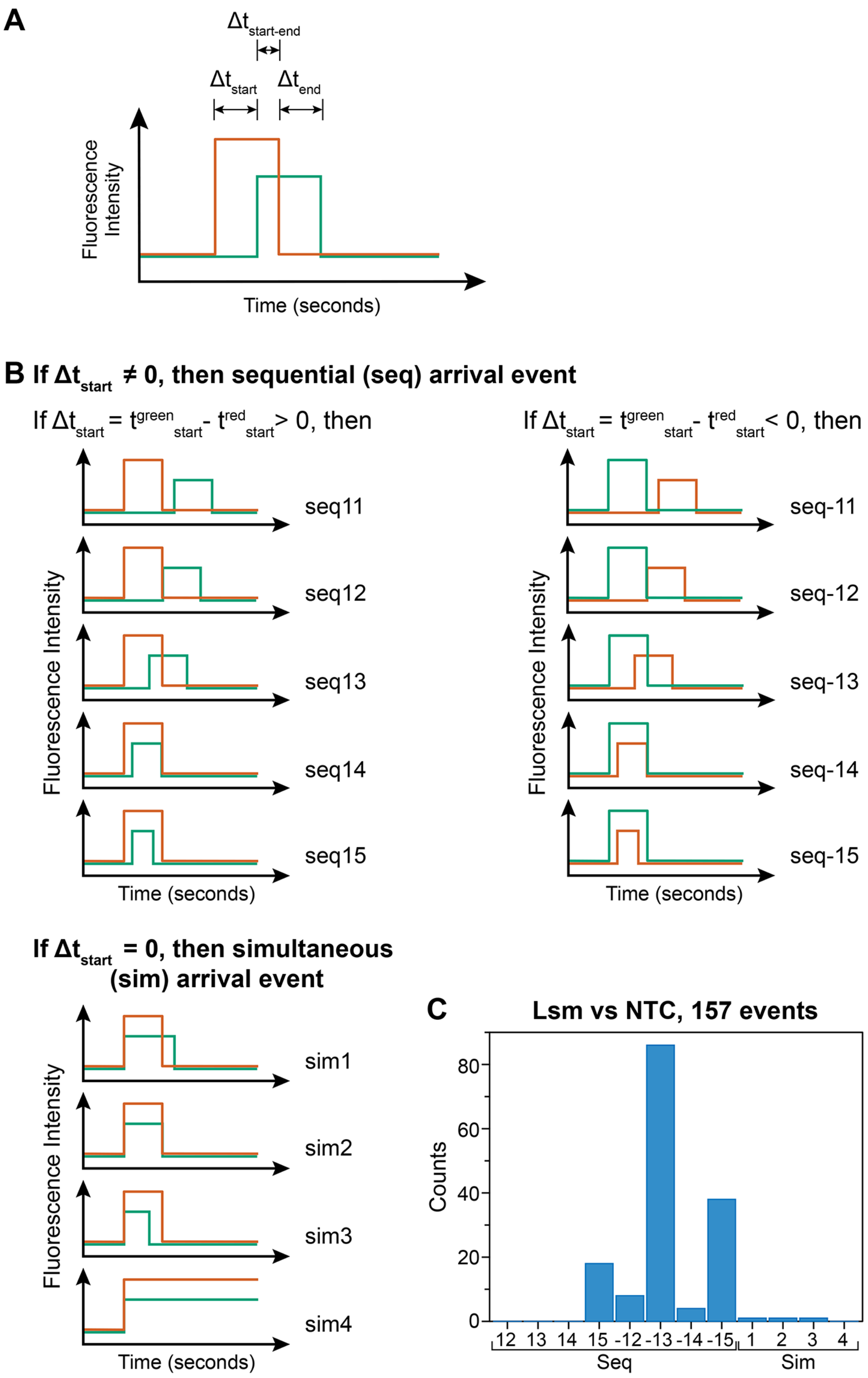


**Figure S8. Scheme for assignment of observed binding and dissociation patterns in CoSMoS assays.** (**A**) For each set of paired events, the beginning and end times of the individual events were recorded. For the analysis shown here, $\Delta t_{start}$ values were first calculated by subtracting the start time of the green event from the start time of the red event. (**B**) $\Delta t_{start}$ values were separated such that all positive values indicated that the red event arrived before the green event (seq11 to 15), negative values indicated that the green event arrived after the red event (seq-11 to -15), and zero values indicated simultaneous arrival (sim1 to 4). Values of $\Delta t_{end}$ and $\Delta t_{start\text{-}end}$ (**A**) were then determined to subcategorize each arrival class based on the disappearance patterns of the green and red events. (**C**) Distribution of Lsm8 and NTC binding events from 3-color CoSMoS experiments. Events of the seq-15 type arise from sampling behaviors.

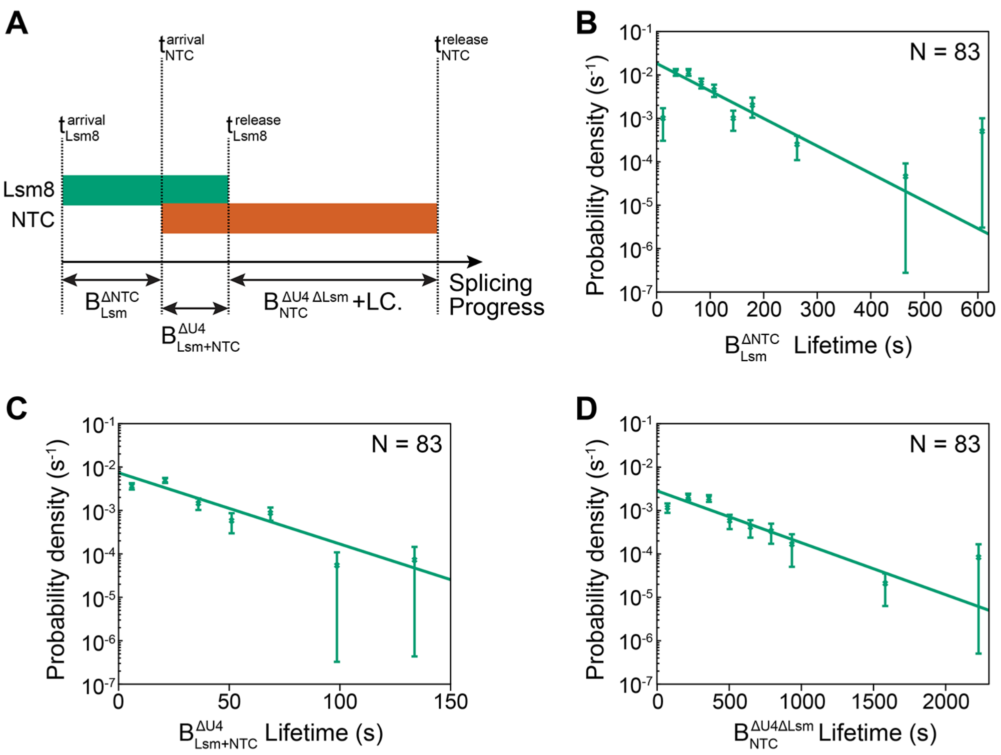


**Figure S9. Lifetimes of Spliceosome Complexes Identified by CoSMoS** (**A**) Schematic showing the relationship between the lifetimes of the $\mathrm{B}_{\mathrm{Lsm}}^{\Delta\mathrm{NTC}}$, $\mathrm{B}_{\mathrm{Lsm+NTC}}^{\Delta\mathrm{U4}}$ , and $\mathrm{B}_{\mathrm{NTC}}^{\Delta\mathrm{U4}\Delta\mathrm{Lsm}}$ and later complexes to the measured arrival and release times for Lsm8 and NTC proteins. (**B-D**) Probability density histograms of $\mathrm{B}_{\mathrm{Lsm}}^{\Delta\mathrm{NTC}}$ (panel B, $\mathrm{t}_{\mathrm{NTC}}^{\mathrm{arrival}} - \mathrm{t}_{\mathrm{Lsm8}}^{\mathrm{arrival}}$), $\mathrm{B}_{\mathrm{Lsm+NTC}}$ (panel C, $\mathrm{t}_{\mathrm{Lsm8}}^{\mathrm{release}} - \mathrm{t}_{\mathrm{NTC}}^{\mathrm{arrival}}$), and $\mathrm{B}_{\mathrm{NTC}}^{\Delta\mathrm{LSM}}$ and later complex lifetimes (panel D, $\mathrm{t}_{\mathrm{NTC}}^{\mathrm{release}} - \mathrm{t}_{\mathrm{Lsm8}}^{\mathrm{release}}$) obtained from the subset of events (*N*) showing ordered arrival of Lsm and then NTC spots followed by ordered loss of the Lsm and then NTC signals. Lines represent fits of the lifetime distributions with equations containing single exponential terms that yielded the parameters reported in **Table S1**. Error bars were calculated for each point as described in the Methods

### Table S1. Fitted Kinetic Parameters

| Tagged protein | Tagged Subcomplex | Complex | Strain | [ATP] mM | $a_1$ | $\tau_1$ (s) | $a_2$ | $\tau_2$ (s) | $a_3$ | $\tau_3$ (s) | Number of Events | Related Figure |
|---|---|---|---|---|---|---|---|---|---|---|---|---|
| Lsm8-SNAP | Lsm2-8 | -- | yAAH1709 | 2 | 0.34±0.03 | 7.5±0.5 | 0.60±0.03 | 69.0±5.0 | 0.06±0.04 | 394.6±116.0 | 700 | Fig. 2 |
| Prp3/Prp4-DHFR | U4 | -- | yAAH1709 | 2 | 0.49±0.10 | 13.8±1.9 | 0.46±0.10 | 41.2±6.7 | 0.06±0.14 | 329.9±72.6 | 481 | Fig. 2 |
| Lsm8-SNAP | Lsm2-8 | -- | yAAH1709 | 0.05 | 0.71±0.05 | 14.2±2.8 | 0.29±0.05 | 543.5±121.3 | NA | NA | 145 | Fig. 2 |
| Prp3/Prp4-DHFR | U4 | -- | yAAH1709 | 0.05 | 0.48±0.05 | 9.0±1.7 | 0.52±0.05 | 413.5±54.8 | NA | NA | 145 | Fig. 2 |
| -- | -- | $\mathrm{B_{Lsm+U4}}$ | yAAH1709 | 2 | NA | 41.7±5.3 | NA | NA | NA | NA | 144 | Fig. 2E |
| -- | -- | $\mathrm{B_{Lsm}^{\Delta U4}}$ | yAAH1709 | 2 | NA | 64.7±6.9 | NA | NA | NA | NA | 144 | Fig. 2F |
| -- | -- | $\mathrm{B_{Lsm}^{\Delta NTC}}$ | yAAH0362 | 2 | NA | 68.6±8.9 | NA | NA | NA | NA | 83 | Fig. S9B |
| -- | -- | $\mathrm{B_{Lsm+NTC}^{\Delta U4}}$ | yAAH0362 | 2 | NA | 21.1±2.7 | NA | NA | NA | NA | 83 | Fig. S9C |
| -- | -- | $\mathrm{B_{NTC}^{\Delta Lsm}}$ | yAAH0362 | 2 | NA | 363.6±38.1 | NA | NA | NA | NA | 83 | Fig. S9D |

**Table S2. Number of Stepping Events Observed During NTC Binding**

| Strain/Subcomplex | Total Events | Step Down Events | Step Up Events |
|---|---|---|---|
| yAAH20 (Syf1/Cef1-SNAP)[a] | 84 | 23 (27%) | 1 (1%) |
| yAAH0362 (Syf1/Cef1-DHFR) (this work) | 83 | 14 (17%) | 0 (0%) |

[a]Data taken from Table S10 (Hoskins et al., 2011).

**Table S3. Oligonucleotide Sequences**

| Oligo # | ID | Sequence (5′-3′) | Length (nts) | Notes |
|---|---|---|---|---|
| 1 | U6_WT | /5/UA CAA AGA GAU UUA UUU CGU UUU /3Phos/ | 23 | WT U6 snRNA (90 - 112nt) |
| 2 | U6_Mut | /5/UA CAA AGA GAU UUA UUU CG**A AAA** /3Phos/ | 23 | Mutate U6 3′ end to polyA |
| 3 | U2_WT | /5/ACG AAU CUC UUU GCC UUU UGG CUU AGA U/3/ | 28 | WT U2 snRNA (1 - 28nt) |
| 4 | U2_Mut | /5/AC**U UUA GAG AAA C**CC UUU UGG CUU AGA U/3/ | 28 | Mutate basepairing region to U6 sequence. |
| 5 | Ah_08_biotinhandle | /5/mAmUmC mCmGmG mAmGmC mGmAmG /iAmMC6T/mAmG mA/3Bio/ | 16 | biotin handle oligo used for coupling to a fluorophore through the internal amino modifier C6T (iAmMC6T) and ligating to RP51A transcripts. |
| 6 | CF13_RP51a_bridge_rnl2 | /5/CTC GCT CCG GAT CGA CCC TTT TGG ATT CTC TTC ATC /3/ | 36 | DNA bridge used for splint ligation of biotin handles to RP51A transcripts, with RNA Ligase II. |

**Table S4. Yeast Strains**

| Strain # | ID | Genotype | Notes |
|---|---|---|---|
| 1 | yAAH0001 | MATa prc1-407 prb1-1122 pep4-3 leu2 trp1 ura3-52 gal2 | BJ2168, parental strain |
| 2 | yAAH0018 | yAAH0001 + cef1::cef1-DHFR-HPH + ntc90::ntc90-DHFR-BLE | contains two DHFR-tagged NTC proteins with hygromycin and phleomycin resistance markers |
| 3 | yAAH0329 | yAAH0001 + prp3::prp3-DHFR-HPH + prp4::prp4-DHFR-BLE | contains two DHFR-tagged U4 snRNP proteins with hygromycin and phleomycin resistance markers |
| 4 | yAAH0362 | yAAH0018 + lsm8::lsm8-$SNAP_f$-NAT | contains two DHFR-tagged NTC proteins and fast-SNAP-tagged lsm8 with nourseothricin resistance marker |
| 5 | yAAH1709 | yAAH0329 + lsm8::lsm8-$SNAP_f$-NAT | contains two DHFR-tagged U4 snRNP proteins and fast-SNAP-tagged lsm8 with nourseothricin resistance marker |
